# Fluid shear magnitude, mechanical history and actin-sensitive coupling shape Piezo1 signalling

**DOI:** 10.64898/2026.10.10.758220

**Authors:** See Swee Tang, Margaret Gheorghita, Benjamin Hansen, Jennifer Frattolin, Konstantinos Kalyviotis, Adele Jing Ying Lam, Simon Tupin, Luana De Abreu Queiros Osorio, Julian Boecker, Joseph van Batenburg-Sherwood, James E. Moore, Victoria Salem, Periklis Pantazis

**Affiliations:** Department of Bioengineering, Imperial College London, London, UK; Centre for Developmental Neurobiology, Institute of Psychiatry, Psychology & Neuroscience, King’s College London, London SE1 1UL, UK; School of Cardiovascular and Metabolic Medicine and Sciences, King’s College, London, UK; School of Engineering, University of British Columbia, Kelowna, BC, Canada; The Francis Crick Institute, London, UK; Randall Centre for Cell and Molecular Biophysics, School of Basic and Medical Sciences, Faculty of Life Sciences and Medicine, King’s College London, London, UK; Making STP, The Francis Crick Institute, London, UK

## Abstract

Mechanical forces shape cell behaviour, yet how Piezo1-associated responses encode force magnitude, recent mechanical history and cytoskeletal context under repeated fluid shear remains unclear. We combined defined fluid shear stress (FSS) with GenEPi, a genetically encoded reporter of Piezo1-proximal Ca²⁺ entry, in engineered XLGenEPi-HEK293T monolayers to resolve mechanically evoked responses at single-cell and population scales. GenEPi responses increased with FSS magnitude but showed diminishing response gain at higher shear. Repeated submaximal FSS produced history-dependent attenuation and reduced temporal coordination, with higher FSS accelerating the early phase of attenuation. Low-dose Latrunculin A, without significantly reducing bulk F-actin abundance, converted sustained FSS-evoked signalling into a brief, decay-dominated response while largely preserving peak responses to Yoda1. These findings show that Piezo1-associated signalling is shaped over minutes by force magnitude, recent mechanical history and actin-sensitive mechanical coupling, providing a quantitative framework for understanding mechanotransduction under sustained and repeated mechanical stimulation.

**Teaser:** Piezo1-proximal imaging links shear magnitude and mechanical history to actin-sensitive mechanical coupling.

## INTRODUCTION

Cells in living tissues are constantly exposed to a broad spectrum of mechanical forces within their microenvironment, including tension, compression and fluid shear stress (FSS) (*1*). These forces shape cellular behaviour, architecture and fate (*2, 3*) and are fundamental to physiological processes including tissue morphogenesis (*4*), immune surveillance (*5*), stem cell differentiation (*5, 6*) and tissue repair (*7*). To respond appropriately, cells rely on integrated mechanisms of mechanosensation and mechanotransduction, through which mechanical forces are detected and converted into intracellular biochemical signals that coordinate adaptive cellular responses.

Mechanosensation and mechanotransduction involve an interconnected network of mechanosensitive membrane proteins, ion channels, adhesion complexes and cytoskeletal structures (*8, 9*). Together, these components maintain cellular mechanical integrity while enabling rapid responses to changes in the physical environment (*10*). Among mechanosensitive ion channels, Piezo1 has emerged as a major mediator of mechanically induced Ca²⁺ entry (*11*). Piezo1 is expressed across a range of mechanically active tissues and contributes to developmental and physiological processes including vascular remodelling (*12*), bone formation (*13*) and lymphatic valve development (*14, 15*). In endothelial cells, Piezo1 responds to FSS generated by blood flow and regulates nitric oxide signalling and cytoskeletal architecture (*16, 17*). Dysregulated Piezo1 signalling has also been implicated in vascular pathologies including atherosclerosis and vascular hyperpermeability (*18, 19*). Recent work has further demonstrated that Piezo1-dependent mechanohydraulic regulation of endocardial cell volume contributes to cardiac morphogenesis through calcium signalling and aquaporin-mediated water transport (*20*).

Piezo1 function can be probed pharmacologically using agonists such as Yoda1 (*21*), while its contribution to mechanically evoked cellular responses is commonly established through genetic or pharmacological perturbation. These approaches establish channel competence or involvement in a mechanical process, but do not necessarily reveal how mechanically evoked Piezo1-associated signalling evolves as the magnitude and history of an applied force change over time. Acute Piezo1 gating and inactivation have been characterised extensively by electrophysiology, with channel sensitivity strongly influenced by membrane tension (*21, 22*), while responses to repetitive mechanical stimulation reveal frequency-dependent filtering of mechanical inputs (*23–25*). These studies have defined Piezo1 behaviour predominantly over millisecond-to-second timescales, including activation, inactivation and recovery from inactivation. By contrast, how Piezo1-associated signalling evolves during sustained and repeatedly applied mechanical stimulation over many minutes remains much less well resolved. Cytosolic Ca²⁺ imaging can follow cellular responses over these longer timescales, but does not distinguish Ca²⁺ entering in the immediate vicinity of Piezo1 from broader intracellular Ca²⁺ signalling. This leaves an important gap between the acute biophysics of Piezo1 gating and the longer-lived channel-proximal signalling responses generated by repeated mechanical stimulation in intact cells. Here, we ask how the magnitude of sustained fluid shear shapes Piezo1-associated signalling across repeated exposures, and whether responses retain a measurable history of previous stimulation across 60-s inter-stimulus intervals that substantially exceed the reported ∼10-s timescale for recovery from Piezo1 inactivation (*23, 24*). A second question concerns the cytoskeletal context of this history-dependent behaviour: although actin is known to influence Piezo1 mechanosensitivity (*22, 25*), whether modest actin perturbation differentially affects sustained mechanical signalling and pharmacological channel responsiveness in intact cells remains unresolved.

Addressing these questions requires experimental platforms that combine mechanical stimulation with channel-proximal measurements over extended recording periods. The combination of a genetically encoded Piezo1-proximal reporter with controlled, sustained fluid shear provides a means to bridge this temporal and methodological gap. We therefore developed a microfluidic imaging platform that applies controlled FSS to engineered XLGenEPi-HEK293T monolayers during live imaging. Building on the genetically encoded GenEPi reporter, which reports Ca²⁺ entry in the immediate vicinity of Piezo1 (*26*), we quantified force-response relationships, repeated-stimulation dynamics, Piezo1-cluster mobility and the effects of mild actin perturbation at single-cell and population scales.

Using this platform, we show that Piezo1-associated signalling is shaped not only by FSS magnitude but also by recent mechanical history and cytoskeletal context. Repeated stimulation produced history-dependent changes in response amplitude and temporal coordination, while low-dose Latrunculin A preferentially impaired sustained mechanically evoked signalling despite largely preserved pharmacological responsiveness. Together, these findings establish a quantitative framework for investigating how Piezo1-associated signalling evolves under dynamic mechanical stimulation.

## RESULTS

### Controlled FSS imaging resolves Piezo1-associated signalling in real time

We established a microfluidic imaging platform that couples controlled fluid shear with real-time imaging of Piezo1-associated signalling using the genetically encoded reporter GenEPi (Fig. 1). Devices were fabricated from polydimethylsiloxane (PDMS) bonded to glass coverslips, functionalised with fibronectin and seeded with XLGenEPi-HEK293T cells (Fig. 1A; Supplementary Materials). Within ∼24 h of seeding, the channels supported uniform adherent monolayers suitable for live imaging (Fig. 1B; fig. S1A,B).

**Figure 1.**
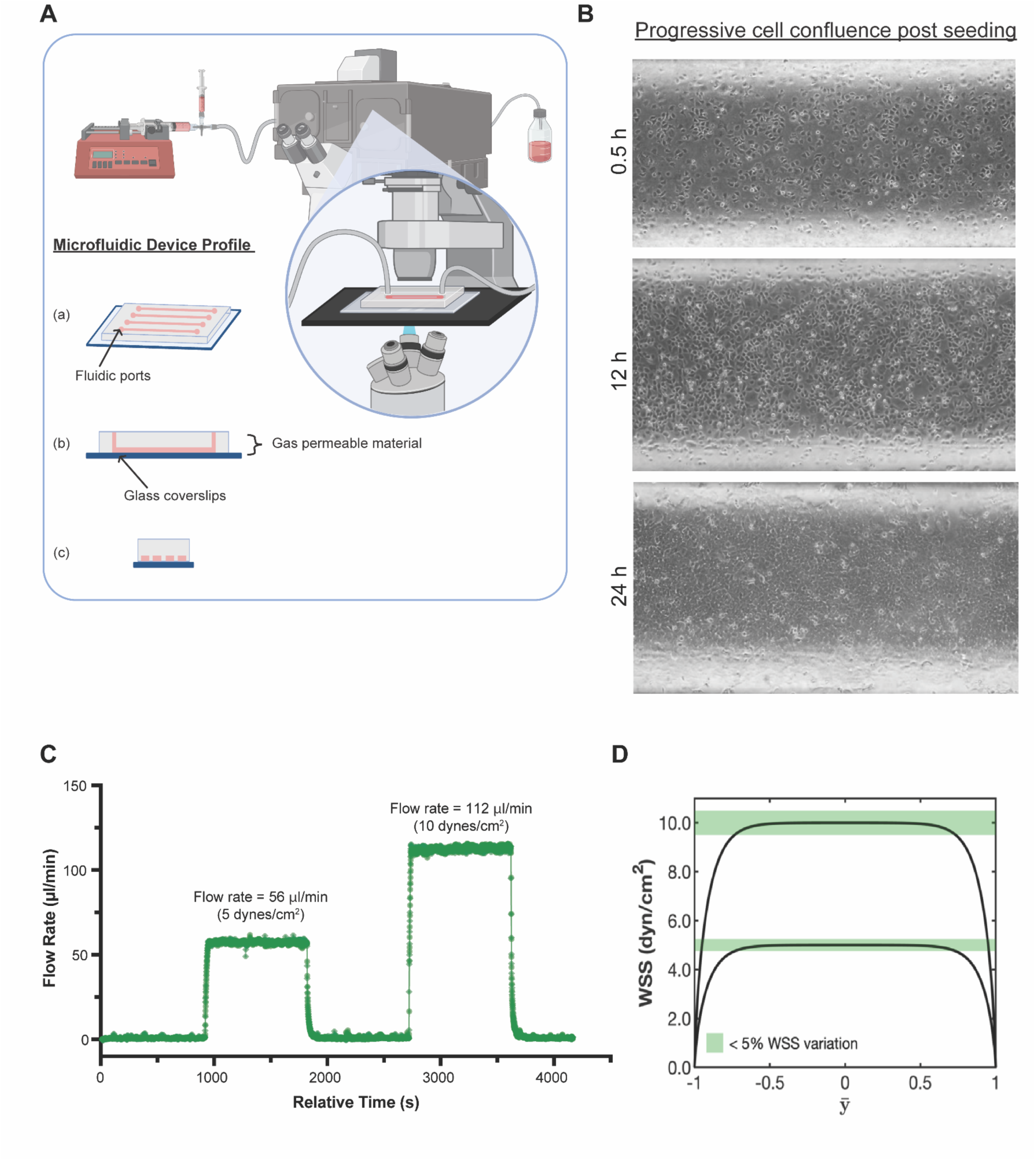
Engineered microfluidic platform enables controlled fluid shear stress delivery during live-cell imaging. (A) Schematic of the microfluidic perfusion and imaging platform. Cells were cultured within a PDMS-based microchannel bonded to a glass coverslip, connected to a programmable syringe pump, and positioned on an inverted microscope for live imaging. Inset: top, side, and cross-sectional views of the device architecture. (B) Representative brightfield images showing progressive formation of an adherent HEK293T monolayer within the microchannel over 24 h. Imaging was performed within this central region. (C) Representative flow-sensor traces during programmed step changes in flow rate. Flow rates of 56 and 112 µl/min, corresponding to nominal wall shear stresses of 5 and 10 dynes/cm², respectively, produced stable plateaus and returned rapidly toward baseline after flow cessation. FSS stimulation was maintained for 1000 s, solely to validate the integrity and stability of flow delivery within our platform, and not used for subsequent biological experiments. (D) Predicted wall shear stress (WSS) distribution across the normalised channel width for nominal 5 and 10 dynes/cm² stimulation. Both profiles contain a broad central region of near-uniform WSS; green shading indicates regions in which WSS varied by <5% from the target value.

To verify delivery of the intended mechanical stimulus, we assembled a syringe-pump-driven flow circuit and monitored the delivered flow profile during imaging (Fig. 1; fig. S1). Step changes in programmed flow produced stable, reproducible plateaus with minimal oscillation and good agreement between programmed and measured flow rates (Fig. 1C). Wall shear stress (WSS) was calculated across the channel width to assess lateral-wall effects (Fig. 1D; fig. S1C). Approximately 73% of the channel width showed <5% deviation from the target WSS, including the central region used for imaging.

We next established an imaging and analysis pipeline to resolve GenEPi dynamics during defined shear. Time-lapse acquisition was aligned to flow onset, and fluorescence was quantified as ΔF/F₀ relative to the pre-flow baseline (Supplementary Materials). Flow onset elicited a rapid increase in GenEPi fluorescence, confirming that mechanically evoked Piezo1-associated Ca²⁺ signals could be resolved during perfusion. GenEPi-positive puncta were segmented and tracked using the ImageJ/Fiji TrackMate plugin (*27*), enabling object-level analysis of fluorescence dynamics and trajectory behaviour under flow.

Together, these data establish a controlled and optically accessible FSS platform for time-resolved measurement of Piezo1-associated signalling, providing the basis for subsequent analyses of force magnitude, mechanical history and GenEPi-positive cluster behaviour.

### Inducible XLGenEPi enables quantitative reporting under flow

To establish a stable reporter system for quantifying Piezo1-associated responses under FSS, we generated a doxycycline-inducible XLGenEPi construct and achieved genomic integration using PiggyBac by co-transfecting XLGenEPi with a CMV-pBase transposase plasmid. Stable HEK293T clones were imaged with or without doxycycline (Dox), and fluorescence was quantified as F/F₀ or ΔF/F₀ (Fig. 2A,B; fig. S2A,B).

**Figure 2:**
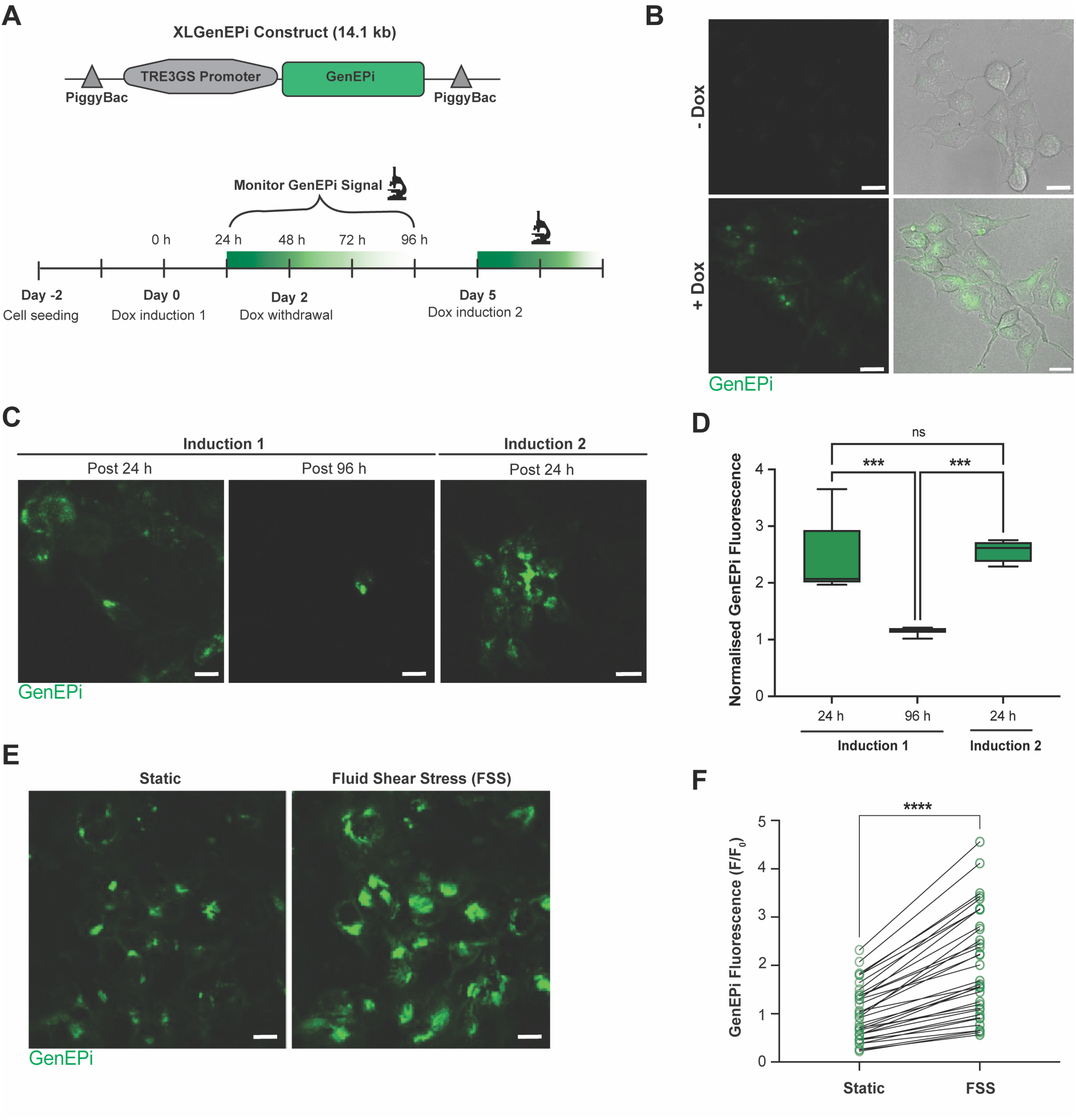
Optical performance and inducible expression of XLGenEPi in microfluidic-cultured HEK cells. (A) Experimental strategy for inducible XLGenEPi expression in cells. Top: Schematic of the 14.1 kb linearised XLGenEPi construct with a TRE3GS promoter driving GenEPi expression, flanked by PiggyBac transposon elements. Bottom: Timeline of doxycycline (Dox)-induced expression. Cells were seeded on Day 2 and treated with two 24 h Dox pulses (Days 0 and 5), separated by withdrawal periods to assess reversibility and re-inducibility. (B) Representative fluorescence images of XLGenEPi-HEK cells in the absence or presence of Dox. Robust GenEPi fluorescence was observed only in the presence of Dox, confirming tight inducibility of expression. Scale bars = 15 µm. (C) Live imaging of GenEPi expression post induction. During Induction 1, strong signal was detected at 24 hours and diminished by 96 hours. During Induction 2, re-exposure to Dox at Day 5 restored GenEPi expression within 24 hours, demonstrating reversible and inducible reporter expression. (D) Quantification of GenEPi fluorescence intensity (a.u.) over time. GenEPi signal increased following the first Dox induction, declined over 96 hours during Dox withdrawal, and was restored following second induction, demonstrating reversible regulation of XLGenEPi expression (*N* = 3 biological repeats, ∼52 cells/time point; One-way analysis of variance (ANOVA); ***, *P* = 0.0005 (Induction 1, 24 h vs. Induction 1, 96 h) and 0.0003 (Induction 1, 96 h vs. Induction 2, 24 h); n.s., not significant). (E) GenEPi response to mechanical stimulation. Representative live-cell images before and after fluid shear stress (FSS) onset showing increased GenEPi fluorescence, indicating Piezo1 activation by flow. (F) Paired single-cell quantification of GenEPi signal before and after 5 dynes/cm^2^ FSS stimulation. Fluorescence intensity (F/F₀) significantly increased after flow application (*N* = 3 biological repeats, ∼34 cells/condition; Wilcoxon test; **** *P* < 0.0001).

Dox induction produced a robust increase in GenEPi fluorescence (∼ 4.86-fold), which declined after withdrawal with a ∼30–32 h half-life and returned to induction levels following re-induction (*N* = 3 biological replicates, ∼52 cells per time point), demonstrating reversible control of reporter expression (Fig. 2C,D).

We next assessed whether XLGenEPi remained pharmacologically responsive in the imaging configuration. The Piezo1 agonist Yoda1 increased GenEPi fluorescence by ∼ 5.03-fold (***, *P* < 0.0001), whereas Ruthenium Red, a non-selective blocker of several Ca²⁺-permeable cation channels, reduced the response towards baseline (ns versus control; N = 3 biological replicates, ∼32 cells per condition; fig. S2C,D). These observations are consistent with preserved Piezo1-linked reporter responsiveness in the microfluidic imaging system.

Finally, laminar FSS elicited a significant increase in GenEPi fluorescence during in-channel imaging (paired F/F₀; *N* = 3 biological replicates, ∼34 cells per condition; ***, *P* < 0.0001), demonstrating that the optical and mechanical configuration was sufficient to resolve stimulus-locked Piezo1-associated responses during perfusion (Fig. 2E,F).

Together, these experiments establish a genomically integrated, inducible XLGenEPi line that retains pharmacological and mechanically evoked responsiveness under flow, providing the quantitative reporter system used for the analyses that follow.

### Piezo1 responses scale with fluid shear stress magnitude and approach a plateau

Having established that inducible XLGenEPi reports mechanically evoked Piezo1-associated calcium signals (Fig. 2), we next examined how these responses depend on FSS magnitude. XLGenEPi-HEK293T cells were exposed to a single laminar FSS cycle consisting of a 60-s baseline followed by 240 s of continuous shear at 1, 2.5, 5 or 10 dynes/cm², selected to span a broad range of low-to-high shear stresses and resolve force-dependent changes in Piezo1-associated signalling, while GenEPi fluorescence was imaged over time (Fig. 3A).

**Figure 3.**
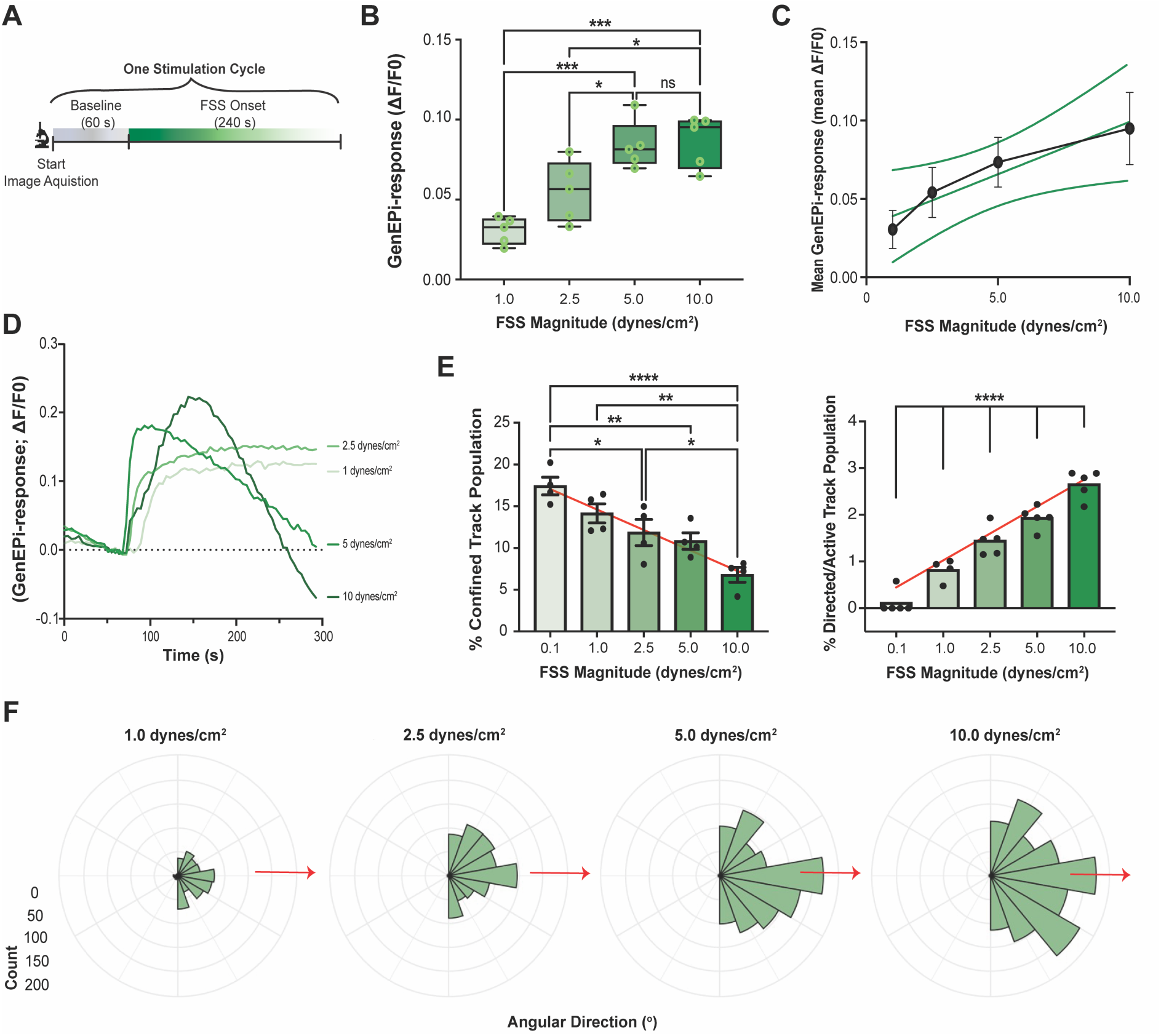
Fluid shear stress elicits graded Piezo1 responses and alters Piezo1-cluster behaviour. (A) Experimental design for single-cycle fluid shear stress (FSS) stimulation. Cells were imaged during a 60-s baseline followed by 240 s exposure to 1, 2.5, 5 or 10 dynes/cm² FSS. (B) Peak cluster-level GenEPi responses from tracked puncta increased with FSS magnitude, with progressively smaller incremental increases at higher FSS (*N* = 5 biological replicates; *n* = 56, 47, 40, 50 and 49 average number of cluster tracks for 0.1, 1, 2.5, 5 and 10 dynes/cm², respectively; One-way analysis of variance (ANOVA); ***, *P* = 0.00080; *, *P* = 0.0236). (C) Replicate-level relationship between applied FSS magnitude and peak GenEPi response. Individual points represent biological replicates and larger symbols show mean ± SEM. Responses increased across the lower-to-intermediate shear range, with little additional increase at higher FSS, consistent with an apparent high-shear plateau. The line connecting condition means is shown as a visual guide only. (D) Population-mean kinetic profiles of GenEPi activation under the different FSS magnitudes. Increasing FSS altered both activation kinetics and the subsequent decay profile during sustained stimulation. (E) Quantification of Piezo1-cluster motility classifications. Increasing FSS decreased the proportion of confined trajectories while increasing the proportion of active trajectories (*N* = 5 biological replicates; *n* = 47, 40, 50 and 49 average no. of tracks for 1, 2.5, 5 and 10 dynes/cm², respectively; One-way analysis of variance (ANOVA); ****, *P* < 0.0001; **, *P* < 0.0080 (0.1 vs. 5.0 dynes/cm^2^) and 0.0032 (0.1 vs. 10 dynes/cm^2^); *, *P* = 0.0236). (F) Polar histograms showing the orientation of Piezo1-cluster displacement during FSS. Trajectory orientations were significantly non-uniform at every FSS magnitude (Rayleigh test; all tests *P* < 0.0001), with a progressive shift in the mean direction of movement toward the flow axis. Mean resultant lengths were similar across FSS conditions.

GenEPi fluorescence increased with increasing FSS magnitude, demonstrating graded sensitivity of Piezo1-associated signalling to the strength of the applied mechanical stimulus (Fig. 3B). Replicate-level analysis showed that peak GenEPi responses increased across the lower-to-intermediate shear range, whereas the increase became smaller at higher FSS (Fig. 3C). In particular, responses showed little additional increase between 5 and 10 dynes/cm², consistent with an apparent high-shear plateau. Thus, across the tested range, Piezo1-associated response amplitude was force dependent but showed diminishing gain at higher mechanical loads.

In addition to the Piezo1-associated response amplitude, mechanical force also altered the temporal characteristics of Piezo1 activation. Higher FSS magnitudes produced distinct activation kinetics, including shorter rise time to half-maximal signal over much of the tested force range, although this trend did not extend to the highest shear level (fig. S3A). Moreover, each FSS magnitude elicited a distinct activation-decay profile: low shear produced slower, more sustained responses, whereas higher shear (≥ 5 dynes/cm²) induced a sharp peak followed by rapid attenuation during continued stimulation (Fig. 3D). Thus, increasing FSS altered not only response magnitude but also the rate at which Piezo1-associated signalling developed and attenuated.

High FSS at 5 and 10 dynes/cm² was also accompanied by modest reductions in nuclear major and minor axes (fig. S3C,D), indicating an early structural response to increased mechanical load. Together, these observations show that increasing FSS magnitude is associated with coordinated changes in both the amplitude and temporal dynamics of Piezo1-associated signalling rather than through a simple on/off response.

To determine whether increasing force also altered the behaviour of individual Piezo1 clusters, we tracked GenEPi-positive puncta during FSS stimulation. Higher FSS progressively reduced the proportion of confined trajectories while increasing the proportion of actively moving clusters (Fig. 3E), indicating that stronger FSS is associated with greater Piezo1 cluster mobility. Moreover, GenEPi-positive cluster trajectories exhibited significant directional organisation at all FSS magnitudes (Rayleigh test, *P* < 0.0001 for each condition; Fig. 3F). Mean resultant lengths remained consistently high across shear conditions (*R* ≈ 0.90-0.92), indicating similarly concentrated trajectory distributions. Increasing FSS progressively shifted the mean direction of Piezo1 cluster movement towards the flow axis, while the overall spread of trajectory directions remained similar across conditions.

### Repeated FSS attenuates Piezo1-associated responses and reduces temporal coordination

Because attenuation at ≥ 5 dynes/cm² developed substantially within a single 240-s exposure, resolving how prior stimulation shapes subsequent responses required lower, submaximal FSS, at which attenuation unfolded over multiple exposures and could therefore be tracked across successive cycles. We therefore examined how Piezo1-associated signalling evolves during repeated stimulation in this lower shear range. Here, XLGenEPi-HEK293T cells were exposed to repeated cycles of submaximal laminar FSS (1 or 2.5 dynes/cm²), separated by 60-s recovery intervals, while GenEPi fluorescence was monitored continuously (Fig. 4A). This experimental paradigm was designed to determine whether repeated exposure to an identical mechanical stimulus alters the magnitude or temporal organisation of Piezo1 activation.

**Figure 4.**
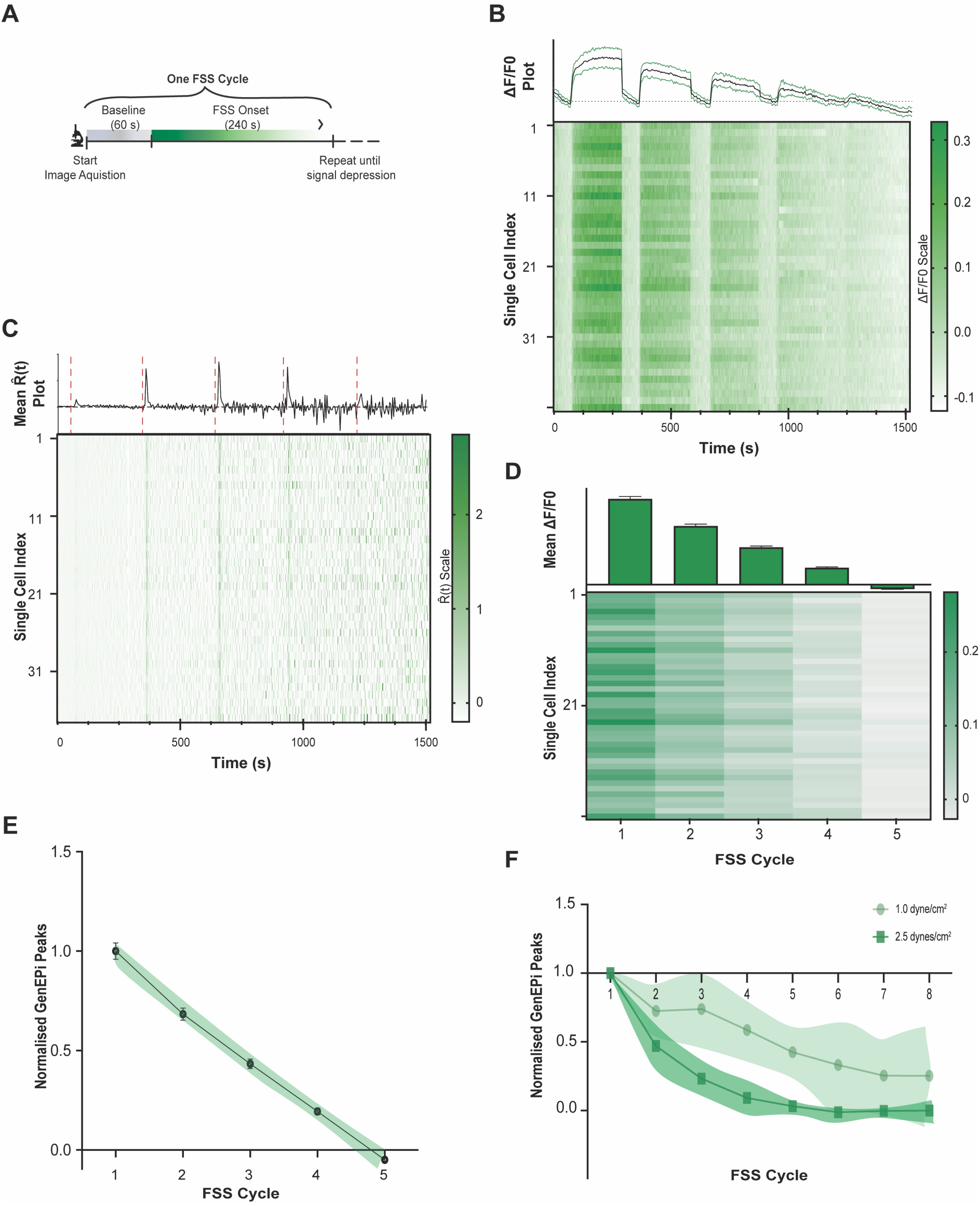
Repeated FSS attenuates Piezo1-associated responses and reduces temporal coordination. (A) Experimental design for repeated FSS stimulation. XLGenEPi-HEK293T cells were imaged continuously during repeated cycles of a 60-s baseline/recovery period followed by 240 s of laminar FSS stimulation. (B) Population-mean GenEPi responses during the first five cycles of repeated 2.5 dynes/cm² FSS from a representative biological replicate. Top: mean ΔF/F₀ ± SEM referenced to the baseline before the first cycle. Bottom: corresponding single-cell heatmap showing progressive reduction in mechanically evoked response amplitude across successive FSS stimulation cycles. (C) Population temporal-derivative analysis of the replicate responses shown in (B). Top: mean R^(t); dashed lines mark FSS onset. Bottom: corresponding single-cell derivative heatmap. Stimulus-associated positive derivative peaks became progressively smaller and less sharply time-locked with repeated stimulation. Because R^(t) is inherently amplitude-sensitive, temporal coordination was assessed independently using dispersion of single-cell response timing (fig. S4C,D). (D) Cycle-resolved peak GenEPi responses for the first five cycles, with corresponding single-cell heatmap. Top: biological-replicate summary. Bottom: corresponding single-cell heatmap illustrating progressive attenuation across the responding population. (E) Peak responses normalised to the first stimulation cycle within each biological replicate; the first five cycles are shown to visualise the early attenuation phase. Data are presented as mean ± SEM; the fitted trend line is shown for visualisation only and was not used for statistical inference. Across the complete eight-cycle 2.5 dynes/cm² series, stimulation cycle had a significant effect on response amplitude (*N* = 3 biological replicates; ∼38 cells/replicate; linear mixed-effects model, *F₇,_18.6_* = 4.09, *P* = 0.0049). (F) Direct comparison of repeated-response trajectories at 1 and 2.5 dynes/cm² FSS. Peak GenEPi responses were normalised to the first stimulation cycle within each biological replicate and are shown as mean ± SEM. Across all eight stimulation cycles common to both conditions, the FSS magnitude × stimulation cycle interaction did not reach statistical significance (*N* = 3 biological replicates per condition; *F₇,_27.06_* = 1.74, *P* = 0.142). Because the 2.5 dynes/cm² response approached baseline by approximately cycle 5, a secondary analysis focused on cycles 1–5 to assess the early attenuation phase before this response reached its floor. This analysis revealed a significant FSS magnitude × stimulation cycle interaction (*N* = 3 biological replicates per condition; *F₄,_16_* = 6.51, *P* = 0.0026), consistent with more rapid early attenuation at 2.5 dynes/cm² than at 1 dyne/cm².

Successive submaximal FSS cycles consistently evoked increases in GenEPi fluorescence. However, the response trajectory differed between shear magnitudes. At 2.5 dynes/cm², response amplitude declined progressively across successive cycles (Fig. 4B). At 1 dyne/cm², by contrast, response amplitude increased over the first few cycles before subsequently declining (fig. S4A). Heatmap visualisation of individual-cell responses showed that attenuation occurred broadly across the responding population rather than arising from a small subset of cells. These observations suggested that the development of history-dependent attenuation may depend on stimulus magnitude, which we tested directly using replicate-level mixed effects analysis.

Because ΔF/F₀ reports GenEPi response amplitude but does not capture the temporal organisation of activation across cells, we computed a temporal derivative-based metric, R^(t), to provide a population-level readout of Piezo1 response dynamics. Each onset of FSS generated a distinct positive R^(t) peak that was closely coupled to stimulus onset during the initial stimulation cycles (Fig. 4C; fig. S4B). With repeated stimulation, these positive derivative peaks became smaller and less sharply stimulus-locked, indicating progressive changes in the temporal organisation of population-wide Piezo1 responses. Because R^(t) is inherently sensitive to response amplitude, temporal coordination was assessed independently using the timing of individual-cell responses. The distributions of individual-cell response times broadened across successive stimulation cycles at both 1 and 2.5 dynes/cm² (fig. S4C,D), indicating increasing heterogeneity in response timing. Consistent with this pattern, replicate-level quantification of within-replicate time-to-peak dispersion revealed a significant effect of stimulation cycle at both FSS magnitudes (2.5 dynes/cm²: linear mixed-effects model, *F_7_,₂₇* = 3.93, *P* = 0.0034; 1 dyne/cm²: linear mixed-effects model, *F_11_,₁₅* = 5.04, *P* = 0.0080; fig. S4C,D). Together, the population-derivative and amplitude-independent timing analyses show that repeated FSS is accompanied by progressively reduced temporal coordination across the responding cell population.

Cycle-by-cycle quantification confirmed a progressive reduction in mean ΔF/F₀ responses throughout the repeated FSS paradigm following peak responses (Fig. 4D), indicating that attenuation tracked cumulative stimulation history rather than occurring as an abrupt loss of responsiveness. Similar history-dependent attenuation was observed at the lower FSS of 1 dyne/cm², although response amplitudes were smaller than at 2.5 dynes/cm² (Fig. 4D; fig. S4E).

Replicate-level mixed-effects analysis confirmed that the cycle-dependent changes in response amplitude were reproducible across independent experiments. At 2.5 dynes/cm², stimulation cycle had a significant effect on peak GenEPi response amplitude (linear mixed-effects model, *F_7_,₂₁* = 7.37, *P* < 0.0001) and the effect remained significant after normalisation of each biological replicate to its first stimulation cycle (*F_7_,₁₈.₆* = 4.09, *P* = 0.0049; Fig. 4E). At 1 dyne/cm², stimulation cycle likewise had a significant effect on peak GenEPi response amplitude (*F_11_,₂₂* = 3.43, *P* = 0.0088; Fig. S4E), which remained significant after within-replicate normalisation to the first stimulation cycle (*F_11_,₁₉.₀* = 6.81, *P* = 0.00023; fig. S4F).

Because responses at 2.5 dynes/cm² approached baseline by approximately cycle 5, whereas responses at 1 dyne/cm² remained above baseline and attenuated more gradually, we distinguished the early attenuation phase from the complete stimulation window shared by both conditions. Across all eight cycles common to 1 and 2.5 dynes/cm², the FSS magnitude × stimulation cycle interaction did not reach statistical significance (*F₇,_27.06_* = 1.74, *P* = 0.142; Fig. 4F), indicating that the complete repeated-response trajectories were not statistically distinguishable over the full shared series. We therefore performed a secondary analysis focused on cycles 1–5, the interval during which the 2.5 dynes/cm² response was still actively approaching its response floor and thus remained informative about attenuation dynamics. This early-phase analysis revealed a significant FSS magnitude × stimulation cycle interaction (*F₄,_16_* = 6.51, *P* = 0.0026; Fig. 4F), consistent with more rapid early attenuation at 2.5 dynes/cm² than at 1 dyne/cm². These results support a force dependence of the early attenuation trajectory rather than a persistent difference across the entire repeated-stimulation series. Together with the pronounced within-stimulus decay observed at 5 and 10 dynes/cm² (Fig. 3D), the data indicate that mechanical load influences the early temporal evolution of Piezo1-associated signalling.

Successive stimulation cycles were separated by 60 s of rest, an interval substantially longer than reported recovery-from-inactivation timescales for Piezo1 measured electrophysiologically (*23, 24*), making canonical fast channel inactivation alone unlikely to account for the cumulative attenuation. Collectively, these findings reveal a history-dependent response state whose early development is modulated by stimulus magnitude rather than simply by the number of preceding stimulation events.

### Low-dose Lat A preferentially impairs mechanically evoked Piezo1 signalling

Our observation that FSS altered both Piezo1 response dynamics and the mobility and directional behaviour of GenEPi-positive clusters suggested that mechanically evoked Piezo1 signalling may be influenced by the cytoskeletal context of the channel. Because the actin cytoskeleton can influence membrane mechanics, protein mobility and force transmission, we therefore asked whether mild actin perturbation would alter mechanically evoked Piezo1 responses. XLGenEPi-HEK293T cells were treated with 0.1 µM Latrunculin A (Lat A), selected to provide a mild actin perturbation while preserving gross cellular architecture, before exposure to repeated 2.5 dynes/cm² FSS. At this functional concentration of 0.1 µM, total phalloidin-labelled F-actin fluorescence was not significantly reduced relative to control (Fig. 5A), whereas higher Lat A concentrations (0.5 and 1.0 µM) produced a dose-dependent reduction in bulk F-actin signal (fig. S5A). Quantitative GLCM-based texture analysis likewise detected no substantial changes in global F-actin texture at 0.1 µM Lat A (two-sided Welch’s t-test with Holm correction; adjusted *P* > 0.729). Despite the absence of detectable gross changes in bulk F-actin abundance or global texture, the temporal profile of mechanically evoked signalling was markedly altered: control cells developed a sustained increase in GenEPi fluorescence during continuous FSS, whereas 0.1 µM Lat A-treated cells showed only a small, transient response that returned towards baseline despite continued shear (Fig. 5B). Thus, mechanically evoked Piezo1-associated signalling is highly sensitive to low-dose Lat A under conditions that do not produce detectable global depletion or large-scale reorganisation of F-actin.

**Figure 5.**
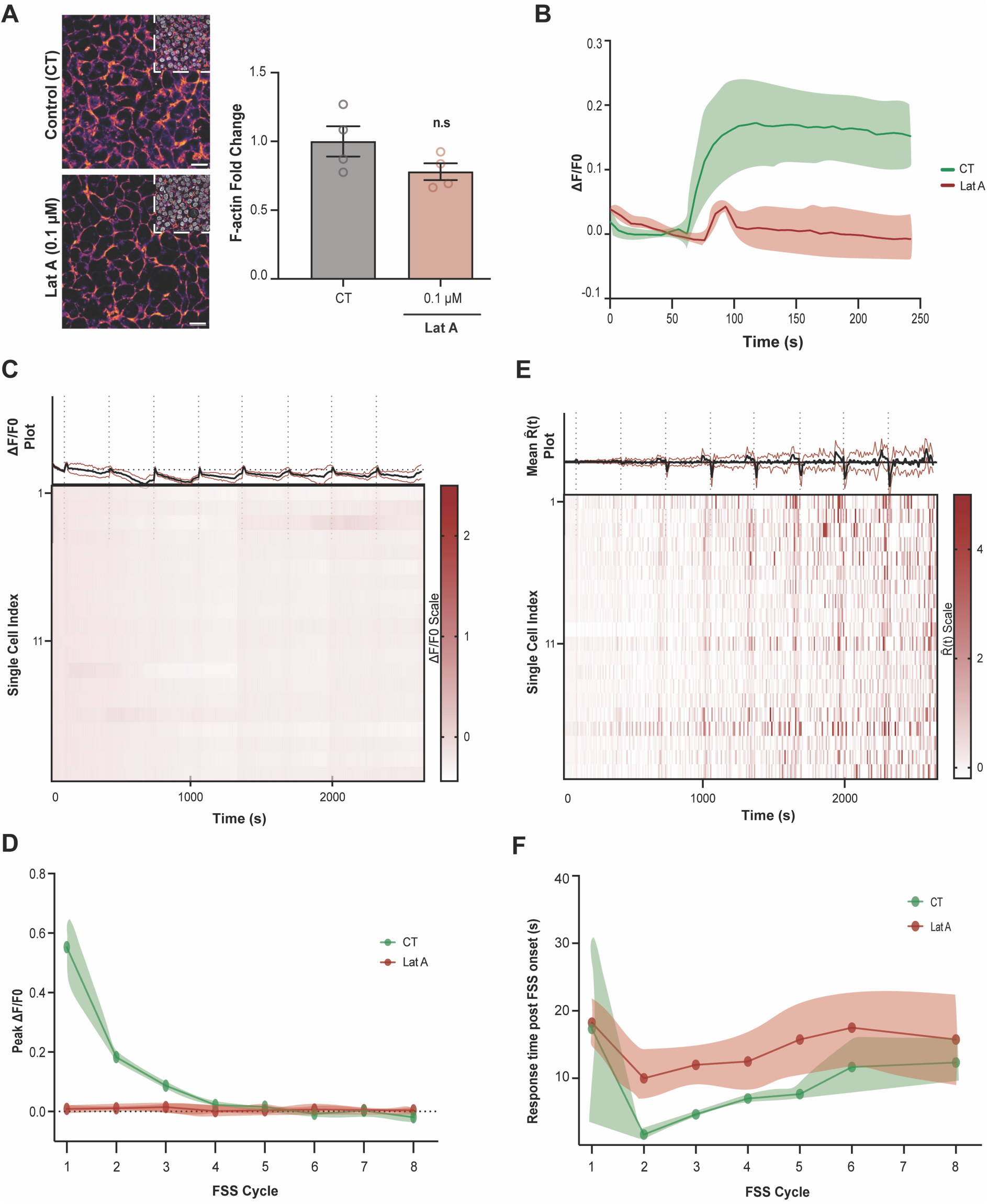
Low-dose actin perturbation selectively impairs mechanically evoked Piezo1 mechanotransduction. (A) Representative phalloidin-labelled F-actin and DAPI-stained nuclei in CT and 0.1 µM Lat A-treated cells. Total F-actin fluorescence was not significantly different at 0.1 µM Lat A, indicating preservation of bulk filamentous-actin abundance at the functional dose (*N* = 3 biological replicates; One-way analysis of variance (ANOVA); n.s, *P* > 0.05). Quantitative GLCM-based texture analysis showed no statistical differences between control and 0.1 µM Lat A-treated cells (*N* = 3 biological replicates; two-sided Welch’s t-test with Holm correction; adjusted *P* > 0.729). Higher-dose titration is shown in fig. S5A. Scale bar = 15 µm. (B) Population-mean ΔF/F₀ trace during the first 2.5 dynes/cm² FSS stimulus in control (CT) and 0.1 µM Lat A-treated XLGenEPi cells. Control cells developed a sustained GenEPi response during continuous FSS, whereas Lat A-treated cells exhibited a markedly smaller, transient response that returned towards baseline despite continued stimulation. Mean responses were interpolated onto a common real-time axis without peak alignment; thick lines show means and shaded regions show ± SEM. (C) Population-wide GenEPi responses during repeated 2.5 dynes/cm² FSS after low-dose Lat A treatment from a representative replicate. Residual stimulus-associated responses remained detectable but were brief and strongly attenuated compared with control responses. (D) Cycle-resolved peak GenEPi responses summarised across independent biological replicates. Peak ΔF/F₀ values extracted from each stimulation cycle remained lower after Lat A during early cycles before converging as responses attenuated, with repeated-cycle responses analysed using linear mixed-effects model (*N* = 3 biological replicates; Linear mixed-effects model; treatment: *F₁,_4_* = 30.26, *P* = 0.0053; stimulation cycle: *F_7_,_28_* = 16.66, *P* < 0.0001; and treatment × stimulation cycle interaction: *F_7_,_28_* = 17.26, *P* < 0.0001). (E) Population temporal-derivative analysis showed smaller and more heterogeneous stimulus-associated responses after Lat A; independent response-time dispersion is used to assess reduced temporal synchronisation (fig. S5E). (F) Response time to the first positive GenEPi derivative response following FSS onset across repeated stimulation cycles in control and low-dose Lat A-treated cells. Response time varied significantly across stimulation cycles (N = 3 biological replicates per condition; linear mixed-effects model; stimulation cycle: F₇,₄₂ = 2.85, P = 0.016), whereas neither the overall treatment effect nor the treatment × stimulation cycle interaction was statistically significant.

Across repeated FSS cycles, control cells showed the progressive attenuation described above, whereas 0.1 µM Lat A-treated cells entered the stimulation series with a markedly blunted response and remained close to baseline across subsequent cycles (Fig. 5C,D; movie S1). Brief stimulus-associated responses remained detectable after Lat A treatment, but these responses rapidly decayed and showed little sustained GenEPi elevation during continued shear. Consequently, the large dynamic range over which control responses progressively attenuated was largely absent after Lat A treatment. Replicate-level mixed-effects analysis confirmed that this altered response trajectory was reproducible across independent experiments. Peak GenEPi responses showed significant effects of Lat A treatment (*F₁,_4_* = 30.26, *P* = 0.0053) and stimulation cycle (*F_7_,_28_* = 16.66, *P* < 0.0001), together with a significant treatment × stimulation cycle interaction (*F_7_,_28_* = 17.26, *P* < 0.0001; Fig. 5D), demonstrating that low-dose Lat A altered both the magnitude and evolution of mechanically evoked Piezo1-associated responses during repeated stimulation.

We next examined how low-dose Lat A altered the temporal dynamics of mechanically evoked Piezo1-associated signalling. In control cells, FSS onset was associated with distinct positive R^(t) transients during early stimulation cycles (fig. S5C). Following Lat A treatment, the initial positive derivative responses were markedly smaller and were frequently followed by negative derivative excursions while FSS was still being applied (Fig. 5E). Because negative R^(t) values denote a population-average decrease in GenEPi fluorescence, these excursions indicate that the mechanically evoked signal decayed after a brief initial activation rather than being maintained throughout the shear stimulus. Consistent with this shift, the GenEPi Response Index moved from activation-dominant values in control cells towards negative values after Lat A treatment (fig. S5D). Thus, low-dose Lat A shifts mechanically evoked Piezo1-associated signalling from sustained activation-dominant dynamics towards a transient, decay-dominated response.

Because R^(t) is amplitude-sensitive, these derivative data do not themselves establish a change in intercellular synchronisation. Temporal coordination was therefore assessed independently using single-cell response-time dispersion. Response-time distributions were broader following Lat A treatment (fig. S5E), and replicate-level mixed-effects analysis revealed significant effects of stimulation cycle (*F_7_,₄₂* = 5.63, *P* < 0.001) and treatment (*F₁,₇* = 9.62, *P* = 0.017) on within-replicate time-to-peak dispersion. However, the treatment × stimulation cycle interaction was not significant (*F_7_,₄₂* = 1.76, *P* = 0.132), indicating that low-dose Lat A was associated with greater temporal heterogeneity overall but did not significantly alter the progressive evolution of temporal coordination across repeated stimulation cycles. Activation response time also varied significantly across the stimulation series (*F_7_,₄₂* = 2.85, *P* = 0.016), whereas neither the overall effect of Lat A treatment nor the treatment × stimulation cycle interaction reached statistical significance (Fig. 5F).

To distinguish impaired mechanical activation from a general loss of Piezo1 responsiveness, we subsequently pharmacologically activated the channel with Yoda1, without any FSS stimulation. Control and Lat A-treated cells exhibited robust concentration-dependent GenEPi responses to 5, 10 and 50 µM Yoda1 (fig. S5F-H). Peak responses were largely preserved after Lat A treatment, whereas response latency was increased at the lower Yoda1 concentrations. Because GenEPi comprises a low-affinity GCaMP tethered to Piezo1 and preferentially reports Piezo1-proximal Ca²⁺ entry, the effects of mechanical and pharmacological stimulation could be compared using the same channel-proximal reporter. The selective loss of sustained FSS-evoked signalling despite largely preserved Yoda1-evoked peak responses therefore argues against a general loss of pharmacological Piezo1 responsiveness and instead points to an actin-sensitive component of mechanical coupling that is particularly important during sustained shear.

Collectively, these data demonstrate that mechanically evoked Piezo1-associated signalling is highly sensitive to mild actin perturbation. Whereas control cells sustained GenEPi elevation during continued shear, low-dose Lat A converted this response into a brief, strongly attenuated transient signal, despite no significant reduction in bulk F-actin abundance. Higher Lat A concentrations produced substantial reductions in F-actin abundance (fig. S5A), confirming concentration-dependent disruption of cytoskeletal integrity. Importantly, substantial Yoda1 responsiveness was retained following low-dose Lat A treatment, indicating that mechanical signalling was preferentially impaired without abolishing pharmacological channel responsiveness. Thus, sustained mechanically evoked Piezo1-associated signalling can be compromised in the absence of detectable bulk F-actin depletion or a general loss of channel responsiveness. Together, these observations are consistent with actin-sensitive cellular mechanics contributing to the maintenance of effective mechanical coupling during sustained shear.

## DISCUSSION

Mechanical forces regulate a broad range of physiological processes, yet how mechanosensitive channels dynamically represent changing mechanical environments remains incompletely understood. Here, controlled microfluidic FSS combined with inducible XLGenEPi imaging revealed that Piezo1-associated responses are shaped by three interconnected features of the mechanical environment: force magnitude, recent stimulation history and actin-sensitive mechanical coupling. Increasing FSS produced graded responses that approached an apparent plateau at higher shear, repeated stimulation progressively attenuated response amplitude and reduced temporal coordination across the responding population, and low-dose Lat A converted sustained shear-evoked signalling into a brief, strongly attenuated response while largely preserving pharmacological activation by Yoda1. Together, these findings support a model in which Piezo1-associated signalling is dynamically shaped by the strength, history and cellular mechanical context of an applied force rather than functioning as a simple binary detector of mechanical stimulation.

Our data show that Piezo1-associated signalling quantitatively reflects FSS magnitude across the tested range. GenEPi responses increased with increasing shear, with diminishing increases at higher mechanical loads and an apparent plateau between 5 and 10 dynes/cm² (Fig. 3). Force magnitude also altered response kinetics: lower shear produced slower and more sustained GenEPi elevation, whereas higher shear generated sharper activation followed by pronounced attenuation during continued stimulation. FSS additionally influenced the spatial behaviour of GenEPi-positive clusters, increasing their mobility and progressively shifting their mean direction of movement towards the flow axis. Dynamic spatial redistribution of Piezo1 has previously been observed in migrating keratinocytes, where local channel enrichment is associated with cellular retraction and migration (*28*). Our findings extend this concept by showing that the mobility and directional organisation of GenEPi-positive clusters are sensitive to the magnitude of an externally applied mechanical load. Thus, increasing mechanical load was associated with coordinated changes in response amplitude, kinetics and Piezo1-cluster behaviour. Together, these findings reveal a multidimensional response that provides a richer representation of mechanical input than a simple on/off activation event.

A second major feature was the dependence of Piezo1-associated signalling on recent mechanical history. Repeated exposure to identical submaximal FSS produced progressive attenuation of GenEPi responses together with increasing dispersion of individual-cell response timing, indicating a progressive reduction in temporal coordination across the responding population (Fig. 4; fig. S4). At 1 dyne/cm², response amplitude initially increased over the first few stimulation cycles before subsequently declining, arguing against simple monotonic fluorescence loss as an explanation for the repeated-stimulation phenotype. Moreover, successive stimuli were separated by 60 s of rest, substantially longer than reported recovery-from-inactivation timescales for Piezo1 measured electrophysiologically (*23, 24*). Canonical fast channel inactivation alone is therefore unlikely to account for the cumulative attenuation observed here. Instead, the response to a given mechanical stimulus depended on the stimulation that preceded it, revealing a history-dependent component of Piezo1-associated signalling that would not be apparent from measurements of a single acute stimulus.

Importantly, force magnitude influenced the early development of attenuation. Across all eight stimulation cycles common to the 1 and 2.5 dynes/cm² conditions, the FSS magnitude × stimulation cycle interaction did not reach statistical significance, as the trajectories converged after the 2.5 dynes/cm² response had approached its floor. By contrast, analysis of cycles 1–5, which captured the phase during which the 2.5 dynes/cm² response was still actively attenuating, revealed a significant interaction, with responses declining more rapidly at 2.5 dynes/cm² than at 1 dyne/cm² (Fig. 4F). Thus, the data support a force dependence of the early attenuation trajectory rather than a persistent difference across the entire repeated-stimulation series. Together with the pronounced within-stimulus decay observed at 5 and 10 dynes/cm² (Fig. 3D), these findings suggest that mechanical load influences how rapidly Piezo1-associated signalling enters an attenuated response state, while later responses converge once attenuation is established.

The actin-perturbation experiments place these dynamic responses in a cytoskeletal context. Our observation that FSS altered both Piezo1 response dynamics and GenEPi-cluster behaviour suggested that mechanically evoked Piezo1 signalling may be influenced by the local cytoskeletal environment of the channel. At 0.1 µM Lat A, total phalloidin-labelled F-actin fluorescence was not significantly reduced relative to control, yet the sustained mechanically evoked GenEPi response was replaced by a small, transient signal that rapidly returned towards baseline despite continued shear (Fig. 5). Higher Lat A concentrations produced concentration-dependent reductions in bulk F-actin fluorescence (fig. S5A), highlighting that the pronounced functional impairment observed at 0.1 µM occurred without detectable global F-actin depletion. Together, these findings suggest that sustained mechanically evoked Piezo1-associated signalling is particularly sensitive to mild actin perturbation, even when bulk F-actin abundance is largely preserved.

The low-dose Lat A condition is particularly informative because Yoda1 continued to evoke robust GenEPi responses, with peak activation largely preserved across the tested concentrations. Because GenEPi comprises a low-affinity GCaMP tethered directly to Piezo1 and specifically reports Piezo1-proximal Ca²⁺ entry, mechanical and pharmacological stimulation could be compared using the same channel-proximal reporter. The selective loss of sustained FSS-evoked signalling despite substantial preservation of Yoda1-evoked peak responses argues against a general loss of Piezo1 responsiveness and instead points to an actin-sensitive step in the coupling of extracellular mechanical load to Piezo1-associated signalling.

This interpretation is particularly interesting in light of previous work showing that the cytoskeleton can influence Piezo1 mechanosensitivity in different ways depending on how force is applied. Removal of cytoskeletal constraints can increase Piezo1 sensitivity to directly applied bilayer tension, consistent with a mechanoprotective role for the cortex under some modes of stimulation (*25*). In intact adherent cells, however, extracellular matrix and cytoskeletal interactions can also shape Piezo1 mechanosensitivity and force transmission (*29—31*). Our findings suggest that the contribution of cortical actin may therefore depend on the geometry and mode of mechanical loading. During fluid shear, membrane-cytoskeletal coupling may help transmit or redistribute mechanical load in a manner that sustains Piezo1-associated signalling, whereas under direct bilayer tension the cortex may instead reduce the force reaching the channel. The persistence of a brief initial response after low-dose Lat A is consistent with some mechanical coupling remaining intact, whereas its rapid decay during continued shear suggests that actin-sensitive cellular mechanics may be particularly important for maintaining effective coupling over time. Notably, quantitative GLCM-based texture analysis likewise detected no substantial alteration in global F-actin organisation at 0.1 µM Lat A treatment, despite the pronounced impairment in mechanically evoked Piezo1 signalling. Together with the absence of a significant reduction in bulk F-actin abundance, these findings suggest that the functional effects of low-dose Lat A are unlikely to arise from large-scale cytoskeletal remodelling. Instead, the pronounced functional effect in the absence of detectable global F-actin changes is consistent with sensitivity to local or dynamic actin-dependent mechanical organisation that is not resolved by bulk fluorescence or global texture measurements.

Taken together, these observations support a model in which force magnitude, mechanical history and cytoskeletal context jointly shape Piezo1-associated signalling (Fig. 6). Force magnitude influences both the initial strength and early temporal evolution of the response. Repeated stimulation progressively reshapes subsequent response amplitude and temporal coordination, and actin-sensitive cellular mechanics influence whether mechanically evoked signalling can be sustained during continued shear. In this framework, higher mechanical load accelerates early entry into an attenuated response state, whereas mild actin perturbation truncates the sustained component of the response from the outset. These findings raise the possibility that the history-dependent state identified here emerges from the cumulative consequences of sustained mechanically evoked signalling rather than simply from repeated channel activation events.

**Figure 6.**
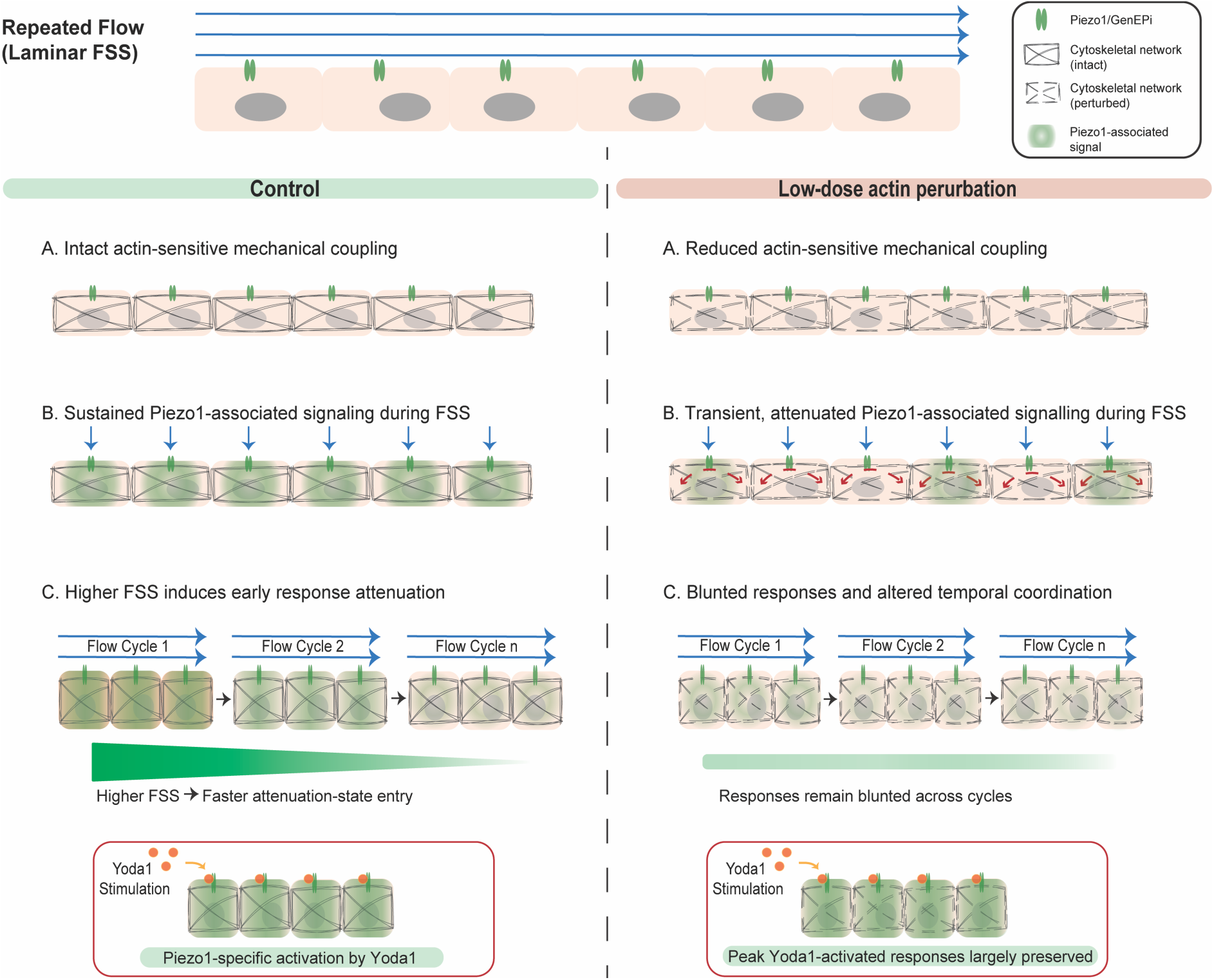
Proposed model for actin-sensitive maintenance of Piezo1-associated responses during repeated FSS. Under repeated laminar fluid shear stress (FSS), control cells exhibit initially stimulus-locked, temporally coordinated Piezo1-associated responses that progressively attenuate and become less temporally coordinated with repeated stimulation. Force magnitude influences the early development of attenuation, with higher FSS accelerating attenuation during the initial stimulation cycles before responses converge towards an attenuated state. Low-dose Latrunculin A (Lat A) converts sustained shear-evoked signalling into a brief, decay-dominated response without significantly reducing bulk F-actin abundance, while largely preserving peak responses to pharmacological activation by Yoda1. The model proposes that actin perturbation compromises the maintenance of mechanically evoked signalling, resulting in persistently blunted responses during repeated stimulation. Together, these findings support a role for actin-sensitive membrane–cytoskeletal mechanics in mechanical coupling to Piezo1-associated signalling, particularly in sustaining responses during continued shear. Green shading represents Piezo1-associated signalling; grey filaments depict intact or perturbed cytoskeletal organisation, and red arrows indicate altered response dynamics following low-dose actin perturbation.

Several limitations nevertheless define important next steps. First, the experiments were performed in an engineered XLGenEPi-HEK293T system that provides precise control over Piezo1 expression, mechanical stimulation and channel-proximal imaging, but does not reproduce the specialised architecture, mechanical environment or signalling networks of native mechanically responsive cells. Determining whether the force-dependent kinetics, history-dependent attenuation and actin-sensitive response persistence described here are conserved in endothelial, epithelial and other Piezo1-expressing cells will therefore be important for establishing their physiological generality.

Second, the repeated FSS paradigm was deliberately simplified, using defined stepwise stimuli to isolate the effects of force magnitude and mechanical history. Cells in vivo instead experience pulsatile, oscillatory and continuously varying mechanical inputs in which force magnitude, duration and recent stimulation history are superimposed. Extending the platform to controlled dynamic waveforms should allow direct testing of how these parameters interact and whether specific temporal features of physiological mechanical inputs are encoded in the amplitude, kinetics and temporal organisation of Piezo1-associated responses.

Finally, the molecular machinery linking Piezo1-associated signalling, history-dependent attenuation and actin-sensitive mechanical coupling remains unresolved. Low-dose Lat A establishes a functional relationship between cytoskeletal context and sustained mechanically evoked signalling, but does not identify the molecular interfaces responsible. Piezo1-centred interactome or proximity-proteomic analyses performed under defined mechanical states could identify candidate components linking the channel to the actin cortex, membrane and adhesion machinery. Targeted perturbation of these candidates could then establish whether they regulate force transmission, sustained Piezo1-associated signalling, history-dependent attenuation or population-level temporal coordination.

In summary, our findings show that Piezo1-associated signalling under fluid shear is shaped jointly by force magnitude, recent mechanical history and actin-sensitive mechanical coupling. The data reveal graded force sensitivity with diminishing response gain at higher shear, history-dependent attenuation accompanied by reduced temporal coordination during repeated stimulation, and a pronounced shift from sustained to transient mechanically evoked signalling following mild actin perturbation, despite largely preserved bulk F-actin abundance. The preservation of substantial Yoda1 responsiveness under low-dose Lat A provides the strongest functional evidence for an actin-sensitive component of mechanical coupling rather than a general loss of Piezo1 responsiveness. More broadly, the findings suggest that force magnitude and mechanical history interact during the early evolution of repeated Piezo1-associated responses, while actin-sensitive cellular mechanics contribute to sustaining the mechanical signal over time. Together with the controlled optical framework established here, this framework provides a quantitative basis for investigating how Piezo1-associated signalling integrates the magnitude, duration and history of dynamic mechanical inputs in physiologically specialised cells and tissues.

## MATERIALS AND METHODS

### Molecular Cloning

Molecular cloning procedure was undertaken to create the XLGenEPi and CMV-pBase constructs, which were used throughout this study. XLone-GFP (*32*) was a gift from Xiaojun Lian (Addgene plasmid #96930; http://n2t.net/addgene:96930; RRID: Addgene_96930). The hGFAPmin-pBase was previously generated in our lab by inserting the hGFAPMIN promoter from pAAV-GFAP-EGFP (a gift from Bryan Roth, Addgene # 50473) into pCAG-PBase plasmid (a gift from Paolo Salomoni). The CMV promoter and GenEPi sensor sequences were derived from the original GenEPi construct (Addgene #140236), previously generated in the Pantazis lab (*26*).

The XLGenEPi and CMV-pBase constructs were generated as follows: the GenEPi sensor sequence was cloned into the XLone plasmid backbone after removal of the EGFP sequence. The pBase sequence from the pCAG-PBase plasmid was cloned into the CMV-backbone of the original GenEPi construct after removal of the sequence.

Following assembly, heat-shock transformation at 37°C was performed using competent E. coli (NEB; K12 strain), plated on LB agar (Thermo Fisher; Q60120 and Q61020), and grown in LB medium (BD Bioscience; 244610) supplemented with 100 µg/ml ampicillin (Sigma-Aldrich, A9393). Bacterial clones were screened by restriction digestion and DNA sequencing (Eurofins Genomics, https://eurofinsgenomics.eu/) using custom primers (table S1). Validated plasmids were isolated using Miniprep (NEB, T2010S) or Maxiprep (Invitrogen; BenchProTM 2100 machine).

### Cell Culture

HEK293T were cultured at 37oC, 5% CO2, in high-glucose DMEM with GlutaMAX (GibcoTM, 10569010), supplemented with 10% FBS (Sigma-Aldrich, F7524) and 1x Penicillin-Streptomycin (Thermo Fisher, 15140122). HEK293T cells used in XLGenEPi co-transfection were at passage 7-8, and XLGenEPi-HEK293T clones were derived from passage 8.

To generate stable XLGenEPi-expressing HEK293T clones, cells were co-transfected with both XLGenEPi and CMV-pBase plasmids. Plasmid DNA amounts were adjusted according to molecular size, since the XLGenEPi construct is several times larger than CMV-pBase and XLone-EGFP. Transfection was performed at 70-80% confluence using Lipofectamine® 3000 (Thermo Fisher, L3000001), following manufacturer’s protocol. Transfection complexes were prepared in Opti-MEM (GibcoTM, 31985062), applied to cells, and incubated overnight (∼16 h) at 37oC, 5% CO2. Cells were refreshed with complete media and allowed to recover for 24 h prior to further use.

For colony selection, cells were dissociated with TrypLETM (GibcoTM, 12604-013) and seeded into 6-well plate (250,000 cells/well). At 70-80% confluence, 10 μg/ml Blasticidin S hydrochloride (BSD; Sigma-Aldrich, 203350) was added for 7 days. Surviving colonies were allowed to recover in fresh media for 3 days. Colonies visible under the EVOS microscope (AMEX1000) at 10x objective were manually picked and transferred by gentle pipetting into 12-well plates for expansion. Each colony was transferred into a separate well. Expanded clones were cryopreserved as XLGenEPi-HEK293T passage 1.

All cultures were routinely tested for mycoplasma contamination (LuBioScience GmBH, B39032). For maintenance, media was changed every two days and cells were passaged at ∼ 70% confluence using TrypLETM. WT HEK293T cells were used between passages 2-8, and XLGenEPi clones between passages 9-10.

### Doxycycline Induction

To induce XLGenEPi expression, cells were cultured in complete growth medium described above, supplemented with 200 ng/mL doxycycline, at least 24 hours prior to seeding into microfluidic devices or glass-bottom dishes for live-imaging.

### Microchannel Device Fabrication and Cell Seeding

A microfluidic system was devised to study Piezo1 dynamics in cell monolayers. Devices were fabricated by PDMS soft lithography, trimmed, and plasma-bonded to No. 1 thickness borosilicate coverglass. Devices were sterilised and coated with fibronectin 1.5 µg/cm2. XLGenEPi-HEK cells were seeded at 10 x 10^6^ cells/ml and allowed to attach before assembling the perfusion circuit. Monolayers were typically established within 24-48 h. Detailed fabrication and seeding protocol are provided in Supplementary Materials.

#### Application of fluid flow

Fluid flow was applied using a PhD Ultra programmable syringe pump (Harvard Apparatus, 70-3007). The required flow rate, *Q*, to achieve a target WSS, τ, along the bottom surface of a rectangular channel was calculated from Eq. 1.

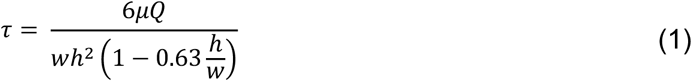

where *μ* is the assumed dynamic viscosity of the perfusion medium (0.93 mPa·s, based on measurements of DMEM supplemented with 10% FBS at 37°C), *h* is the channel height (120 μm), and *w* is the channel width (800 μm). Target WSS values were 0.1, 1, 2.5, 5 and 10 dynes/cm²; withdrawal mode was used for culture perfusion (0.1 dyne/cm²) and infusion mode for shear-stimulation experiments (1–10 dynes/cm²).

To quantify lateral wall effects, the WSS distribution across the channel width was calculated using Eq. 2. The <5% variation region shown in Fig. 1D and fig. S1C was derived from this cross-channel calculation using the same measured channel geometry and viscosity.

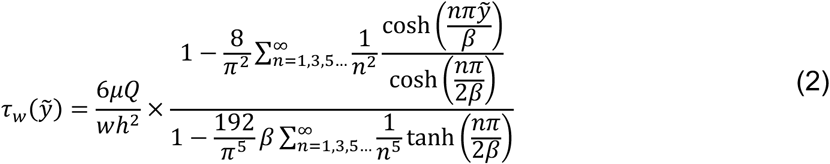

Where *β* = ℎ/*w* and *y*2 = 2*y*/*w*, where *h* is the height of the channel, *w* is the width of the channel, and *y* is the distance to the centre of the chip’s width.

Cells were exposed to laminar FSS using the microfluidic platform described above. For single-stimulation experiments (Fig. 3), cells were imaged for 60 s before application of continuous FSS for 240 s at the indicated shear magnitude. For repeated-stimulation experiments (Figs. 4 and 5), each cycle consisted of a 60-s baseline/recovery period followed by 240 s of continuous FSS. Continuous imaging was maintained throughout each repeated-stimulation dataset. Repeated FSS experiments were conducted with a total of 8 FSS stimulation cycles for 2.5 dynes/cm^2^ (both control and Lat A experiments) and 12 FSS stimulation cycles for 1 dynes/cm^2^.

### Pharmacological and Cytoskeletal Perturbations

For the validation of XLGenEPi activation function, cells were stimulated with Yoda1 (10 µM) and then Ruthenium Red (RR; 10 µM) during the imaging period.

To investigate the contribution of the actin cytoskeleton to Piezo1 mechanotransduction without inducing gross cytoskeletal collapse, XLGenEPi-HEK293T cells were treated with 0.1 µM Lat A within the microfluidic device for 5 min before imaging. This low concentration was selected to preserve cell adhesion and viability and, in the main functional condition, did not significantly reduce bulk phalloidin-labelled F-actin fluorescence (Fig. 5A). Parallel titration with 0.5 and 1.0 µM Lat A produced dose-dependent reductions in F-actin fluorescence (fig. S5A). Immediately before imaging, Lat A-containing medium was washed out and replaced with fresh culture medium, after which live-cell imaging was performed on cells subjected to either the indicated FSS stimulation protocols or pharmacological activation by Yoda1.

For pharmacological activation of Piezo1 during live-imaging, cells were treated with Yoda1 at final concentration of 5, 10, or 50 µM. Live-cell imaging was performed 500 s, including a 60 s pre-treatment baseline followed by Yoda1 application.

### Brightfield and Confocal Imaging

Brightfield images were acquired on an EVOS XL Core microscope (Thermo Fisher, AMEX1000) under 10x magnification to monitor seeding and cellularisation.

Confocal imaging was performed on a Leica SP8 microscope equipped with a white light laser using an HC PL APO CS2 20x/0.75 dry objective. For live imaging, cells were excited at 488 nm for GenEPi signal and acquired at 8-bit, 512 x 512 resolution in photon-counting mode. Regions of interest were imaged across 20 µm z-plane depth over experimental duration using bidirectional scanning with offset correction for speed. Experiments were acquired at the temporal sampling interval range of 4.24-5.39 s per frame, depending on the z-stack sizes. Live imaging was carried out in Live Cell Imaging Solution (Thermo Fisher, A14291DJ) or culture medium, pre-warmed and equilibrated to 37°C before imaging, with stage-top incubation at 37°C. Live Cell Imaging solution was used for static GenEPi imaging, while culture medium was used for all the FSS experiments.

For chemical perturbations, 10 µM Yoda1 (Tocris Bioscience, 5586) or 10 μM Ruthenium Red (Sigma-Aldrich, R2751; in water) were added directly during imaging.

Fixed-cell imaging was performed at 8-bit, 1024 x 1024 resolution, with excitation at 405 nm (DAPI), 488 nm (Alexa Fluor 488), and 647 nm (Phalloidin). Z-stack images were acquired at 0.88 µm steps across 20 µm depth.

### Immunofluorescence Staining

Cells in microchannels were fixed with 4% paraformaldehyde (Thermo Fisher, 28908) for 10 min at 37oC, permeabilized with DPBS containing 0.1% TritonX-100 (Sigma-Aldrich, T8787), and blocked with 10% donkey serum (Jackson ImmunoResearch, 17-000-121). Primary antibodies were diluted in 5% donkey serum/0.1% TritonX-100, followed by fluorophore-conjugated secondary antibodies and DAPI (Thermo Fisher, 62248). Following final washes, channels were filled with DPBS and stored in 4oC until imaging. Antibodies used are listed in table S2.

### Image Processing and Quantification

ImageJ/Fiji (v1.53f51) (*33*) was used for image analysis. For immunofluorescence, Z-stacks maximum-projected. For live imaging, GenEPi (Piezo1) clusters were segmented from 8-bit images by background subtraction (rolling ball radius of 2 pixels), median filtering (radius of 2 pixels), and Otsu thresholding, followed by watershed and dilation. Quantitative features (e.g., mean gray value) were extracted using “Analyze Particles”.

#### *GenEPi fluorescence processing and ΔF/F*₀ *quantification*

Time-lapse fluorescence images were processed using ImageJ/Fiji (v1.53f51). Custom image and statistical analyses scripts were written and performed within RStudio Integrated Development Environment (v2026.04.0+526; RStudio Team, 2026) with R Statistical Software (v4.6.1; R Core Team, 2026).

Individual GenEPi-positive puncta or regions of interest were identified following background subtraction and intensity thresholding as described above. Fluorescence intensity was measured for each tracked object at every time point and expressed as the fractional change in fluorescence relative to the pre-stimulation baseline:

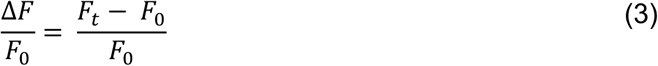

where *F_t_* is the fluorescence intensity at time *t* and *F_0_* is the mean baseline fluorescence measured before stimulation.

For single-FSS experiments, baseline fluorescence was calculated from the 60-s pre-stimulation period immediately preceding flow onset. For repeated FSS, continuous image acquisition was maintained throughout the stimulation protocol, and displayed ΔF/F₀ traces were calculated relative to the baseline preceding the first stimulation cycle to preserve a common reference across the recording. Individual cycles were segmented according to the programmed FSS onset times. Cycle-resolved peak responses were extracted separately for quantitative comparison, as described below.

Biological replicates acquired at different temporal sampling intervals were linearly interpolated onto a common real-time grid before calculation of condition-level summary traces. Importantly, interpolation was performed solely to align sample times between biological replicates; fluorescence traces were not time-shifted, response-aligned, or peak-aligned. Mean responses and associated variability (standard deviation or standard error of mean, as specified in the corresponding figure legends) were calculated independently at each interpolated time point.

Quantitative response metrics, including peak ΔF/F₀, response timing and cycle-resolved response amplitudes, were extracted from individual biological replicates prior to statistical analysis. Where smoothing was applied to displayed mean traces, it was used exclusively for visualisation and not for quantitative measurements or statistical testing.

#### Temporal derivative analysis of population-wide Piezo1 activity

To describe the temporal organisation of mechanically evoked GenEPi responses across the recorded population, a temporal-derivative analysis was performed (*34, 35*). For each analysed cell, the normalised fluorescence response was calculated as:

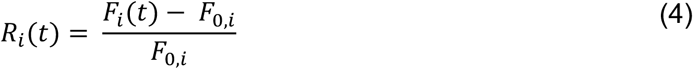

where *F_i_(t)* is the fluorescence intensity of cell *i* at time *t*, and *F_0,i_* is the corresponding pre-stimulation baseline fluorescence.

To characterise rapid temporal changes in GenEPi activity, the first temporal derivative of each response trace was calculated. To reduce slow baseline offsets between traces, the time-averaged derivative of each individual recording was subtracted, yielding a centred temporal derivative:

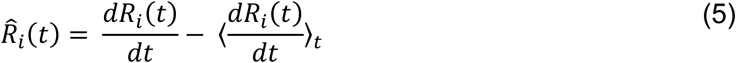

where 〈. 〉*_t_* denotes the temporal mean over the analysed recording.

Population-level temporal response profiles were obtained by averaging the centred derivatives across all analysed cells at each time point,

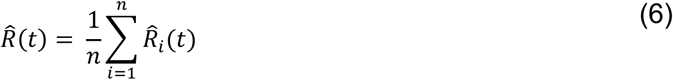

where n is the total number of analysed cells. Positive values indicate population-average increases in GenEPi fluorescence and negative values indicate population-average decreases. Distinct positive peaks therefore identify stimulus-associated activation dynamics. Broader or attenuated derivative peaks may reflect changes in the temporal coordination of population-wide responses; however, because the derivative magnitude depends on response amplitude and signal-to-noise ratio, this metric cannot independently establish cellular temporal coordination.

To determine the GenEPi Response Index, quantitative analysis was performed within the first 120 s following each FSS onset. A maximum positive derivative and minimum negative derivative were extracted for each cycle, and the GenEPi Response Index was calculated as:

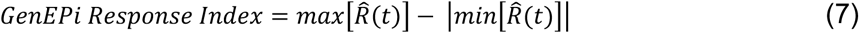

where positive values indicate activation-dominant responses and negative values indicate depression-dominant responses. Response latency was defined as the elapsed time between FSS onset and the first positive peak in *R*^4^(*t*) within the analysis window.

#### Peak response and response timing analysis

For each analysed cell, the peak mechanically evoked GenEPi response was defined as the maximum ΔF/F₀ value, observed within the stimulation period following the FSS onset. For repeated FSS experiments, peak GenEP responses were extracted independently from each stimulation cycle to enable cycle-resolved quantification of response amplitude. Mean peak responses were calculated for each biological replicate prior to statistical analysis.

Where indicated, continuous fluorescence traces from independent biological replicates acquired at different temporal sampling intervals were linearly interpolated onto a common real-time axis before averaging. Interpolation was performed solely to align sampling times between biological replicates; fluorescence traces were not time-shifted, response-aligned, peak-aligned, or otherwise normalised in time.

For experiments examining repeated FSS stimulation, cycle-resolved peak ΔF/F₀ responses were quantified throughout the stimulation series to assess changes in mechanically-evoked Piezo1 activity. Where indicated, peak responses were normalised to the corresponding first stimulation cycle within each biological replicate to facilitate comparison of response attenuation between independent experiments. Representative cycle-resolved response profiles were generated from individual biological replicates for visualisation, whereas quantitative analyses were performed using cycle-resolved peak responses extracted independently from each biological replicate.

To directly compare repeated-response trajectories between 1 and 2.5 dynes/cm² FSS, cycle-resolved peak responses were normalised to the first stimulation cycle within each biological replicate. A primary linear mixed-effects model was fitted across the eight stimulation cycles common to both FSS conditions, with FSS magnitude, stimulation cycle and their interaction specified as fixed categorical effects and biological replicate included as a random intercept. Because responses at 2.5 dynes/cm² approached baseline by approximately cycle 5, later cycles primarily reflected persistence near the response floor rather than continued attenuation. To specifically assess the early attenuation phase, a secondary model using the same fixed and random effects was therefore fitted over cycles 1–5. *F*-tests used Satterthwaite approximations for denominator degrees of freedom.

Response latency was defined as the elapsed time between FSS onset and the first positive peak in the corresponding population temporal-derivative trace within the predefined analysis window. For pharmacological activation experiments, response latency was measured as the interval between Yoda1 addition and the first positive peak in the corresponding temporal-derivative trace. Peak response amplitudes and response latencies were calculated independently for each biological replicate before statistical analysis.

#### Single-cell temporal coordination analysis

To assess the temporal coordination independent of GenEPi response amplitude, a single-cell response timing metric was quantified directly from original, non-interpolated fluorescence recordings. Repeated FSS recordings were automatically segmented into individual complete stimulation cycles according to the experimental protocol (60 s baseline followed by 240 s FSS). For each cell and stimulation cycle, the local baseline fluorescence was calculated from the 60 s preceding FSS onset. Cells were classified as responders when the maximum ΔF/F₀ within the first 120 s after the FSS onset exceeded the baseline mean by > 3 standard deviations. Response timing was defined as the elapsed time between FSS onset and the peak ΔF/F₀ response. Non-responding cells were retained for calculation of responder fractions but excluded from time-to-peak analyses.

Temporal coordination was quantified independently for each biological replicate as the within-replicate standard deviation of single-cell time-to-peak values for each stimulation cycle, with interquartile range calculated as a robustness measure where indicated. For control versus Lat A comparisons, replicate-level timing dispersion was analysed using linear mixed-effects models with treatment, stimulation cycle and their interaction as fixed effects and biological replicate as a random effect. Cycle-specific comparisons were performed using estimated marginal means with Holm correction for multiple testing.

#### Piezo1 cluster tracking and trajectory analysis

Individual GenEPi-positive Piezo1 clusters were tracked using the Fiji/ImageJ plugin, TrackMate (*27*). Cluster centroids were identified from the segmented binary masks described above and linked between consecutive frames using the Linear Assignment Problem (LAP) tracker. Trajectories with insufficient duration or incomplete tracking were excluded from subsequent analyses.

Cluster displacement and trajectory coordinates were exported for quantitative analysis in MATLAB (R2021b) using the @msdanalyser package (*36*). Mean squared displacement (MSD) was calculated for each trajectory to quantify cluster mobility throughout FSS stimulation. Based on the calculated diffusion behaviour, trajectories were classified as either confined or active, and the proportion of clusters within each category was determined for each experimental condition.

To examine the directional behaviour of Piezo1 clusters under FSS, the angular displacement of individual cluster trajectories was calculated relative to the direction of fluid flow and visualised using circular histograms (rose plots). Directionality within each dataset was assessed using the Rayleigh test for circular uniformity. Mean resultant length (*R*; range 0-1) was calculated as a measure of angular concentration, with values approaching 1 indicating greater concentration of trajectories around the mean direction. Circular variance (1 - *R*) and circular standard deviation were additionally calculated to characterise angular dispersion. Mean trajectory direction was determined for each FSS condition relative to the experimentally defined flow axis.

#### Nuclear morphometric analysis

Confocal z-stacks of DAPI-stained nuclei were acquired before and after FSS stimulation using identical imaging settings within each experiment. Maximum-intensity projections were generated in ImageJ/Fiji (v1.53f51), and nuclei were segmented following background subtraction and intensity thresholding. Segmentation masks were visually inspected and corrected where necessary before quantitative analysis.

Nuclear morphometric features (area, perimeter, axes, aspect ratio) were extracted and quantified. The parameters were subsequently compared between experimental conditions to assess structural changes associated with FSS.

#### F-actin fluorescence and texture quantification

Bulk F-actin abundance was assessed from phalloidin-stained confocal images using ImageJ/Fiji (v1.53f51). Following thresholding, regions of interest corresponding to individual cells were defined, and integrated fluorescence density and cell area were measured. To account for differences in cell size, F-actin abundance was expressed as normalised integrated density, calculated as:

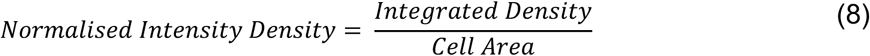

For each biological replicate, normalised integrated density values were subsequently normalised to the mean value of the corresponding untreated control (CT) group to obtain fold-change values for quantitative comparison between control and Lat A-treated groups,

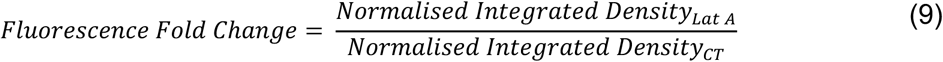

Fold-change values were used for graphical presentation and statistical analysis.

To determine whether low-dose Lat A treatment altered global actin organisation drastically independently of bulk F-actin abundance, quantitative texture analysis was performed on the same phalloidin images using a custom Python script (Python 3.14; scikit-image (*37*)). Original ImageJ composite TIFF image files were analysed using the phalloidin fluorescence channel only. Images were analysed on their original 8-bit intensity scale without any intensity normalisation or histogram stretching. Grey-level co-occurrence matrices (GLCMs) were computed after fixed quantisation to 32 grey levels using pixel offsets of 1, 2 and 4 pixels in four directions (0°, 45°, 90°, and 135°). Standard Haralick texture features (*38*), including contrast, homogeneity, correlation, energy and Shannon entropy, were calculated for each image and averaged across all distances and directions to generate a single texture measurement per biological replicate. These measurements were used to assess whether Lat A treatment altered global F-actin texture under conditions in which bulk F-actin abundance was largely preserved.

To summarise the multidimensional actin texture measurement, all five standardised GLCM-derived texture features were subjected to principal component analysis (PCA). Because the first principal component (PC1) explained the majority of the total variance and received similar contribution from GLCM factors, including contrast, homogeneity, energy, and entropy, it was used as a summary measure of global F-actin texture for graphical presentation and statistical comparison between treatment groups.

### Statistical Analysis

All statistical analyses were performed using GraphPad Prism (v11.1.0(218), GraphPad Software), RStudio Integrated Development Environment (v2026.04.0+526; RStudio Team, 2026) with R Statistical Software (v4.6.1; R Core Team, 2026), and MATLAB (R2021b; MathWorks), as appropriate. The specific statistical tests applied, along with the numbers of analyzed nuclei, regions of interest (ROIs), embryos, and independent experiments, are indicated in the corresponding results and figure legends.

Biological replicates (*N*) represent independent experiments performed on separate culture batches. Individual cells, nuclei, GenEPi-positive clusters or trajectories (*n*) were treated as measurements within a biological replicate and were summarised at the replicate level prior to statistical testing unless otherwise stated. Statistical comparisons were therefore performed using biological replicate values to minimise pseudoreplication.

Comparisons between two independent groups were performed using unpaired two-tailed Student’s t-tests, with Welch’s correction when variances were unequal. Paired comparisons used paired tests where appropriate. Comparisons involving more than two groups were analysed using one-way ANOVA with appropriate multiple-comparison testing, or non-parametric alternatives when distributional assumptions were not met.

For analyses of Piezo1 cluster directionality, circular statistics were performed in MATLAB (*39*). Directional organisation within each experimental condition was evaluated using Rayleigh test for circular uniformity. Mean resultant length (R), circular variance and circular standard deviation were calculated to quantify the degree of directional alignment of cluster trajectories relative to the direction of fluid flow.

For repeated FSS experiments, cycle-resolved peak ΔF/F₀ responses were analysed using linear mixed-effects models (lmer, lme4 package in RStudio (*40*)). For experiments examining repeated FSS in single experimental condition, stimulation cycle was specified as a fixed categorical effect and biological replicate included as a random intercept. For experiments comparing control and Lat A-treated cells, treatment condition, stimulation cycle, and their interaction were specified as fixed effects, with biological replicate included as a random intercept. Independent biological replicates were treated as the experimental unit for all mixed-effects analyses. Where appropriate, post-hoc comparisons between individual stimulation cycles and Cycle 1 were performed using estimated marginal means (emmeans) (*41*) with Holm correction for multiple testing. Adjusted pairwise comparisons were used to annotate significance between experimental conditions or between stimulation cycles where indicated in the figures.

Statistical significance was defined as *p* < 0.05. For all statistics, p-value numbers were reported in text and graphically depicted as n.s. = *p* > 0.05, * = *p* < 0.05, ** = *p* < 0.01, *** = *p* < 0.001 and **** = *p* < 0.0001 in the results.

## Supporting information

Supplementary Movie S1: Piezo1-associated responses to repeated fluid shear stress

## Acknowledgments

We thank members of the Pantazis laboratory for helpful discussions. We also thank Leica Microsystems for their support through the Imperial College London and Leica Microsystems Imaging Hub.

## Author contributions

Conceptualisation: See Swee Tang, Periklis Pantazis.

Formal analysis: See Swee Tang, Periklis Pantazis.

Investigation: See Swee Tang, Margaret Gheorghita, Benjamin Hansen, Adele Jing Ying Lam, Jennifer Frattolin, Simon Tupin, Luana De Abreu Queiros Osorio, Konstantinos Kalyviotis, Julian Boecker.

Methodology: See Swee Tang, Adele Jing Ying Lam, Simon Tupin, Konstantinos Kalyviotis, Julian Boecker.

Project administration: Periklis Pantazis.

Resources: Benjamin Hansen, Adele Jing Ying Lam, Simon Tupin, Luana De Abreu Queiros Osorio, James E. Moore Jr, Joseph van Batenburg-Sherwood, Victoria Salem, Periklis Pantazis. Supervision: Jennifer Frattolin, James E. Moore Jr, Joseph van Batenburg-Sherwood, Victoria Salem, Periklis Pantazis.

Validation: See Swee Tang, Margaret Gheorghita, Benjamin Hansen, Periklis Pantazis.

Visualisation: See Swee Tang, Margaret Gheorghita, Jennifer Frattolin, Periklis Pantazis. Funding acquisition: Periklis Pantazis.

Writing - original draft: See Swee Tang, Periklis Pantazis.

Writing - review & editing: See Swee Tang, Margaret Gheorghita, Benjamin Hansen, Adele Jing Ying Lam, Jennifer Frattolin, Simon Tupin, Luana De Abreu Queiros Osorio, Konstantinos Kalyviotis, Julian Boecker, James E. Moore Jr, Joseph van Batenburg-Sherwood, Victoria Salem, Periklis Pantazis.

## Supplementary Materials and Methods

### PDMS-based straight-channel device

Design and master fabrication. A mould (fig. S1) was designed in Inventor 2022 (Autodesk, USA) and printed on an Objet 500 3D printer (Stratasys, USA) using VeroWhite (Stratasys, USA). This master was used to cast the PDMS devices used throughout the study. The device-production and cell-seeding workflow was as follows:

(A) Device fabrication. PDMS (Sylgard 184, Dow) was mixed at 10:1 (base:curing agent), degassed under vacuum, and 5 g of the mixture was poured over the 3D-printed mould. Polymerisation was performed at 80 °C for ≥4 h (typically overnight). The cured PDMS was trimmed, 1-mm inlet/outlet ports were created with a biopsy punch (Kai Medical, BPP-10F), and the PDMS slab was plasma-bonded to a #1 borosilicate coverslip (24 × 50 mm; VWR 631-0146) using a Piezobrush PZ2 (Relyon Plasma), followed by baking at 80 °C for 5 min to strengthen the bond.

(B) Device preparation and coating. Devices were sterilised by autoclaving (121 °C, 15 min), dried at 70 °C, and wetted by rinsing ≥3× with 1× DPBS (Sigma-Aldrich, D8537). Channels were coated with fibronectin (Sigma-Aldrich, F0895) or 0.1% gelatin (Sigma-Aldrich, G9391) as indicated (fig. S1A).

For fibronectin coating, a 1 mg/ml stock was diluted on ice in 1× DPBS containing Ca²⁺/Mg²⁺ (Gibco, 14040083) to achieve 1.5 µg/cm² on the channel glass. Coating solution was pipetted through each channel from both ports at least twice, with ∼30 µl left at each port to minimise bubble ingress. Devices were incubated in a humidified chamber at 37 °C and 5% CO₂ overnight.

(C) Device seeding. HEPES-supplemented culture medium was prepared the day before seeding by adding 10 mM HEPES (Gibco, 15630080) to complete DMEM, filtering through a 0.22-µm membrane, and equilibrating overnight at 37 °C and 5% CO₂. Following coating, channels were rinsed with DPBS containing Ca²⁺/Mg²⁺ and equilibrated with HEPES-supplemented medium. XLGenEPi-HEK293T cells were detached with TrypLE (Gibco, 12604-013; 5 min, 37 °C), resuspended to 10 × 10⁶ cells/ml, and introduced into the devices. For HEK293T cells, viability within the channels was supported by an additional 10% FBS. Fluids were introduced from the designated inlet to maintain one-way flow. Devices were returned to the incubator for ≥20 min to allow attachment, the seeding step was repeated once, and devices were incubated for ∼4 h before connection to the flow circuit.

(D) Microchannel cell culture. After attachment was confirmed, 15 µl of pre-warmed HEPES-supplemented medium (with the additional FBS used for XLGenEPi-HEK293T cells) was gently perfused through the inlet. The flow circuit was assembled as shown in fig. S1. After equilibration, the outlet syringe was mounted on the pump and flow was initiated at 1.12 µl/min for 24 h, corresponding to 0.1 dynes/cm² in the rectangular channel.

**Figure S1.**
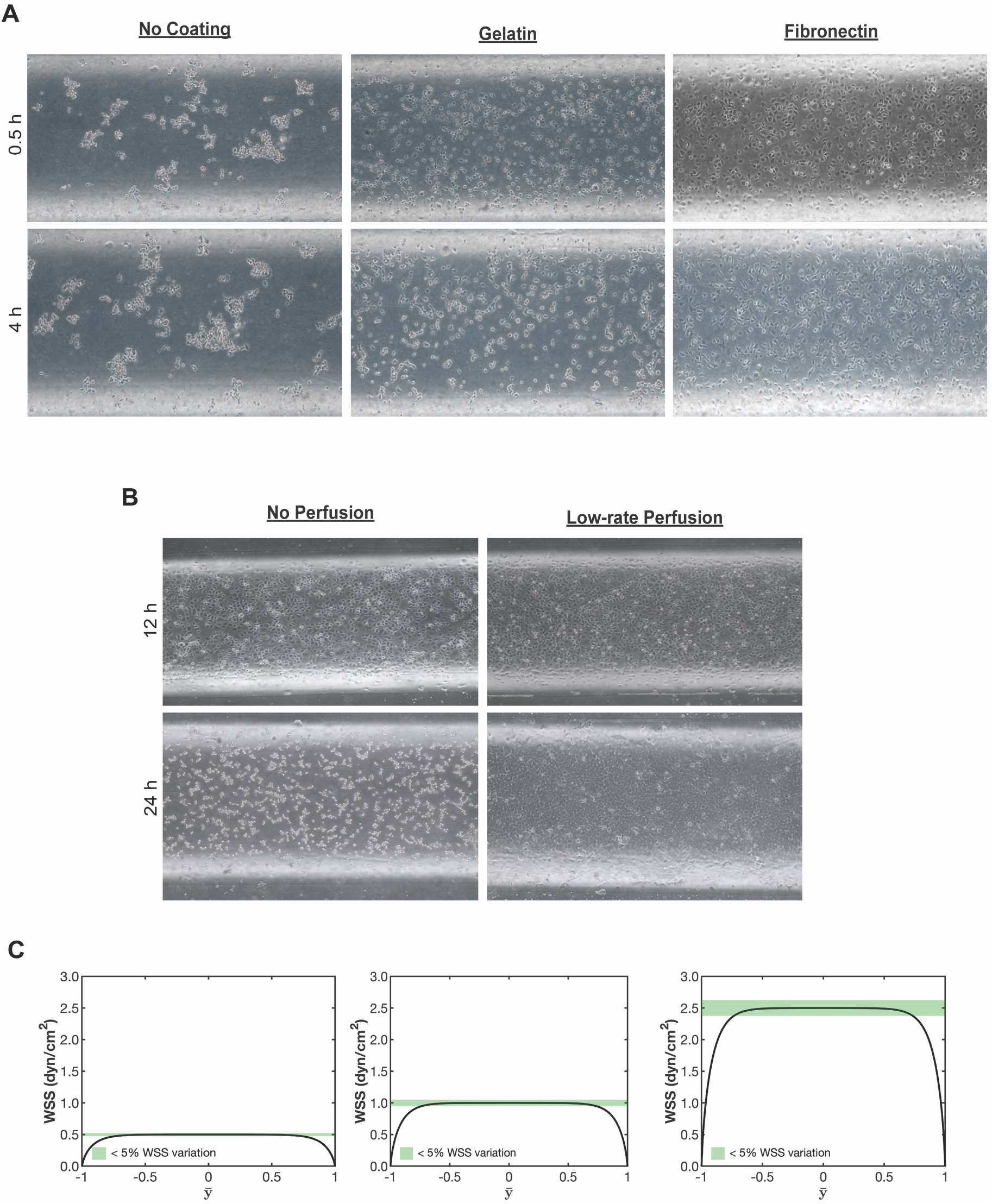
Optimisation and validation of the microfluidic culture platform. (A) Bright-field images at 0.5 and 4 h after seeding under the indicated coating conditions. Fibronectin coating provided the strongest cell attachment and retention within the microfluidic channel. (B) Comparison of static culture and continuous low-rate perfusion after 12 and 24 h, showing improved cell retention during perfusion. (C) Predicted WSS profiles for nominal 0.5, 1, and 2.5 dynes/cm²; green shading denotes <5% deviation from the target WSS. The 0.5 dynes/cm² profile is shown for device characterisation purpose and not used in subsequent biological experiments.

**Figure S2.**
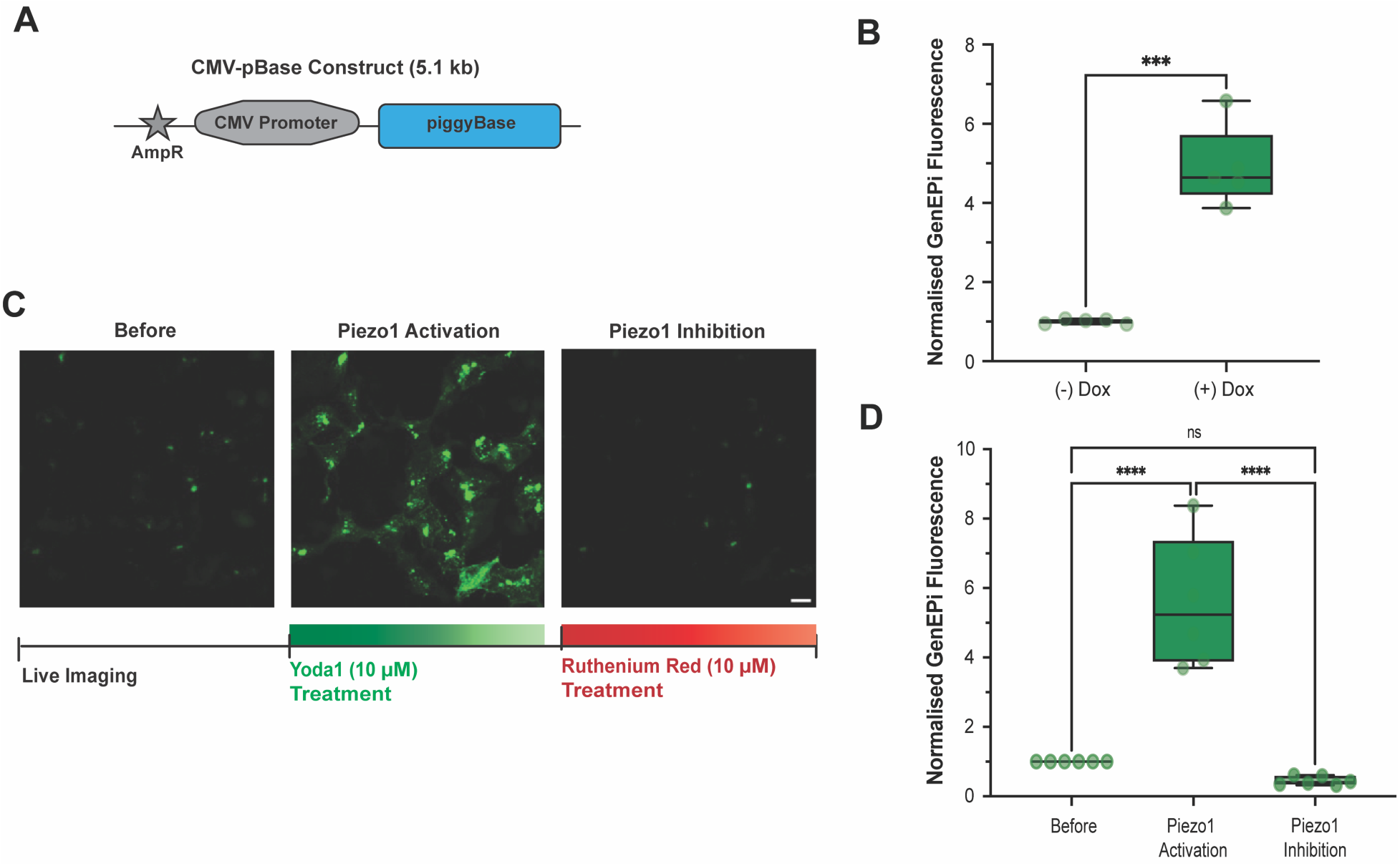
Stable genomic integration and pharmacological validation of XLGenEPi responsiveness. (A) Schematic of the 5.1-kb linearised CMV-pBase plasmid encoding piggyBac transposase. CMV-pBase was co-transfected with the XLGenEPi construct to mediate stable genomic integration of the reporter cassette. (B) Normalised baseline GenEPi fluorescence in XLGenEPi-HEK293T cells cultured without or with doxycycline. Doxycycline induced robust reporter expression (*N* = 4 biological repeats; ∼35 cells per condition; Welch’s *t* test; *** *P* = 0.0010). (C) Representative live-cell images before treatment, after Yoda1 (10 µM), and after subsequent Ruthenium Red (10 µM). Scale bar, 15 µm. (D) Quantification of GenEPi fluorescence. Yoda1 increased the signal by ∼5-fold, whereas subsequent Ruthenium Red reduced the signal toward/below baseline (*N* = 3 biological repeats; ∼32 cells per condition; One-way analysis of variance (ANOVA); **** *P* < 0.0001; n.s., not significant).

**Figure S3.**
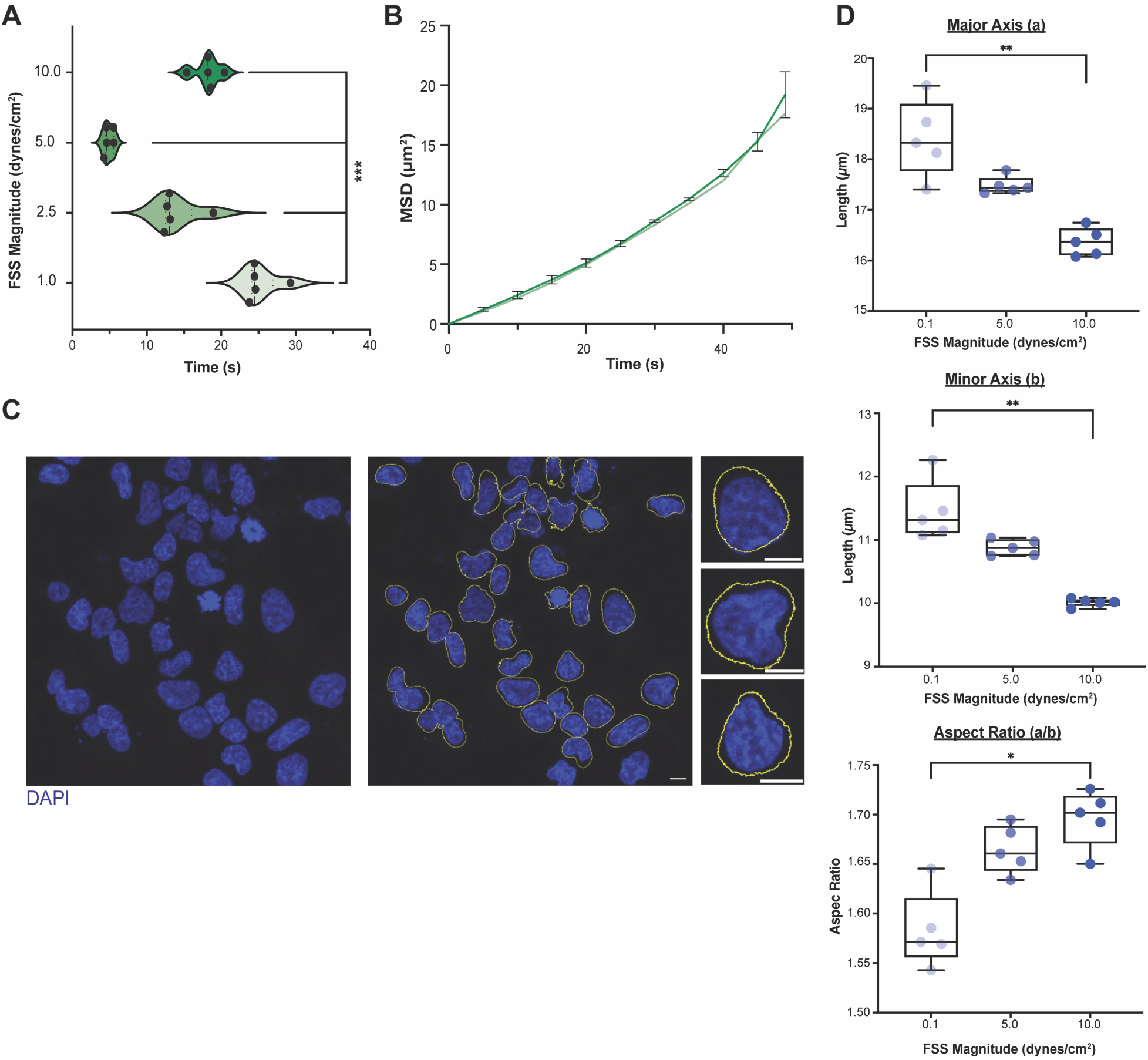
Shear-dependent Piezo1 dynamics and nuclear morphology. (A) Time-to-half-maximal GenEPi response across single-cycle FSS stimulation at 1, 2.5, 5, and 10 dynes/cm². Activation kinetics varied with FSS magnitude, with the shortest response time observed at 5 dynes/cm², whereas 10 dynes/cm² did not further accelerate response onset (*N* = 5 biological replicates; *n* = 47, 40, 50, and 49 tracks for 1, 2.5, 5, and 10 dynes/cm², respectively; One-way analysis of variance (ANOVA); ***, *P* = 0.00020). (B) Mean-square-displacement (MSD) analysis of GenEPi-positive cluster trajectories during FSS stimulation. MSD increased throughout stimulation, indicating continued cluster movement under FSS. Data are presented as mean ± SEM. (C) Representative DAPI images acquired before and after high-shear exposure, with segmented nuclear boundaries indicated. Scale bars, 15 µm (main images) and 5 µm (insets). (D) Nuclear major axis, minor axis, and aspect ratio across the indicated FSS conditions. Increasing FSS was associated with modest reductions in major and minor nuclear axes, whereas aspect ratio was comparatively preserved. (*N* = 5 biological replicates; *n* = 33, 45, and 39 average no. of nuclei for 0.1, 5, and 10 dynes/cm², respectively; One-way analysis of variance (ANOVA); *, *P* = 0.0133; **, *P* = 0.0047 for indicated comparisons). Individual nuclei are shown for visualisation.

**Figure S4.**
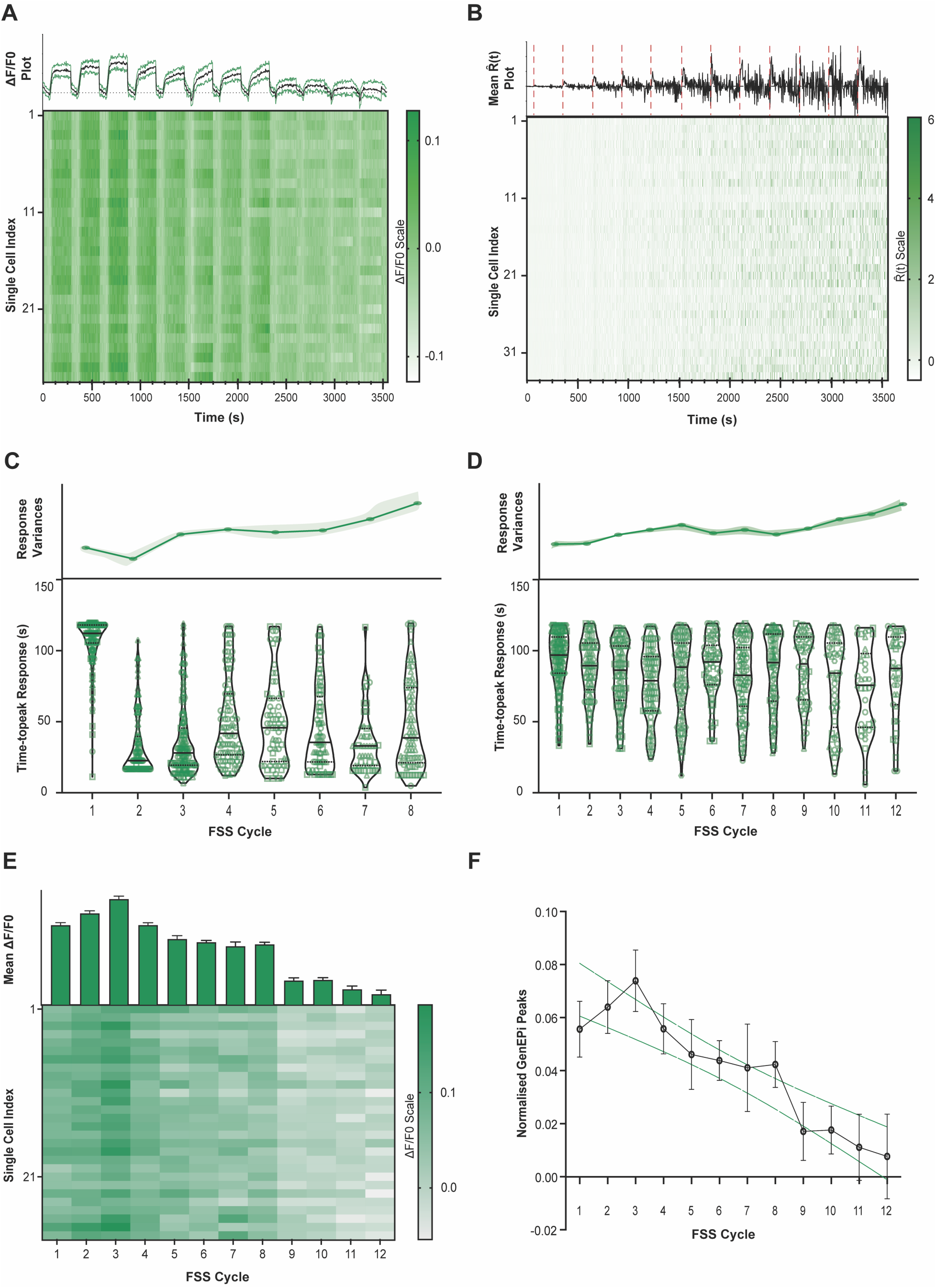
Repeated low-magnitude FSS reveals history-dependent attenuation and progressive broadening of Piezo1 responses. (A) Population-wide GenEPi responses during repeated 1.0 dynes/cm² FSS from representative biological replicate. Top: mean ΔF/F₀ trace across the analysed population. Bottom: corresponding single-cell ΔF/F₀ heatmap. Responses remained detectable throughout the series, with an early non-monotonic phase followed by attenuation during later cycles. (B) Temporal-derivative representation of the responses shown in (A). Distinct stimulus-associated derivative features were most apparent during early cycles and became less prominent during continued stimulation. This analysis is used as a descriptor of stimulus-locked temporal dynamics and is not an amplitude-independent measure of cell-cell synchrony. (C and D) Dispersion of single cell response timing during repeated FSS stimulation at 2.5 dynes/cm^2^ (C) and 1 dynes/cm^2^ (D). Distribution of individual-cell response times progressive broadened across successive stimulation cycles, indicating increasing heterogeneity in Piezo1 activation timing. Replicate-level mixed-effects analysis confirmed a significant effect of stimulation cycle on within-replicate response-time dispersion (*N* = 3 biological replicates; Linear mixed-effects model; C: 2.5 dynes/cm², ∼38 cells/replicate, *F_7_,₂₇* = 3.93, *P* = 0.0034; D: 1 dyne/cm², ∼43 cells/replicate, *F_11_,₁₅* = 5.04, *P* = 0.0080). (E) Cycle-resolved peak GenEPi responses across independent biological replicates, with the corresponding single-cell heatmap. Peak response increased over the first few cycles before declining across later stimulation cycles; this non-monotonic behaviour is inconsistent with a simple monotonic loss of fluorescence through photobleaching alone (*N* = 3 biological replicates; ∼43 cells/replicate; Linear mixed-effects model; *F_11_,_22_* = 3.43, *P* = 0.0088). (F) Peak responses normalised to the first stimulation cycle within each biological replicate to facilitate comparison of attenuation between experiments. After within-replicate normalization, stimulation cycle remained a significant determinant of response amplitude (*N* = 3 biological replicates; Linear mixed-effects model; *F_11_,₁_9.0_* = 6.81, *P* = 0.00023).

**Figure S5.**
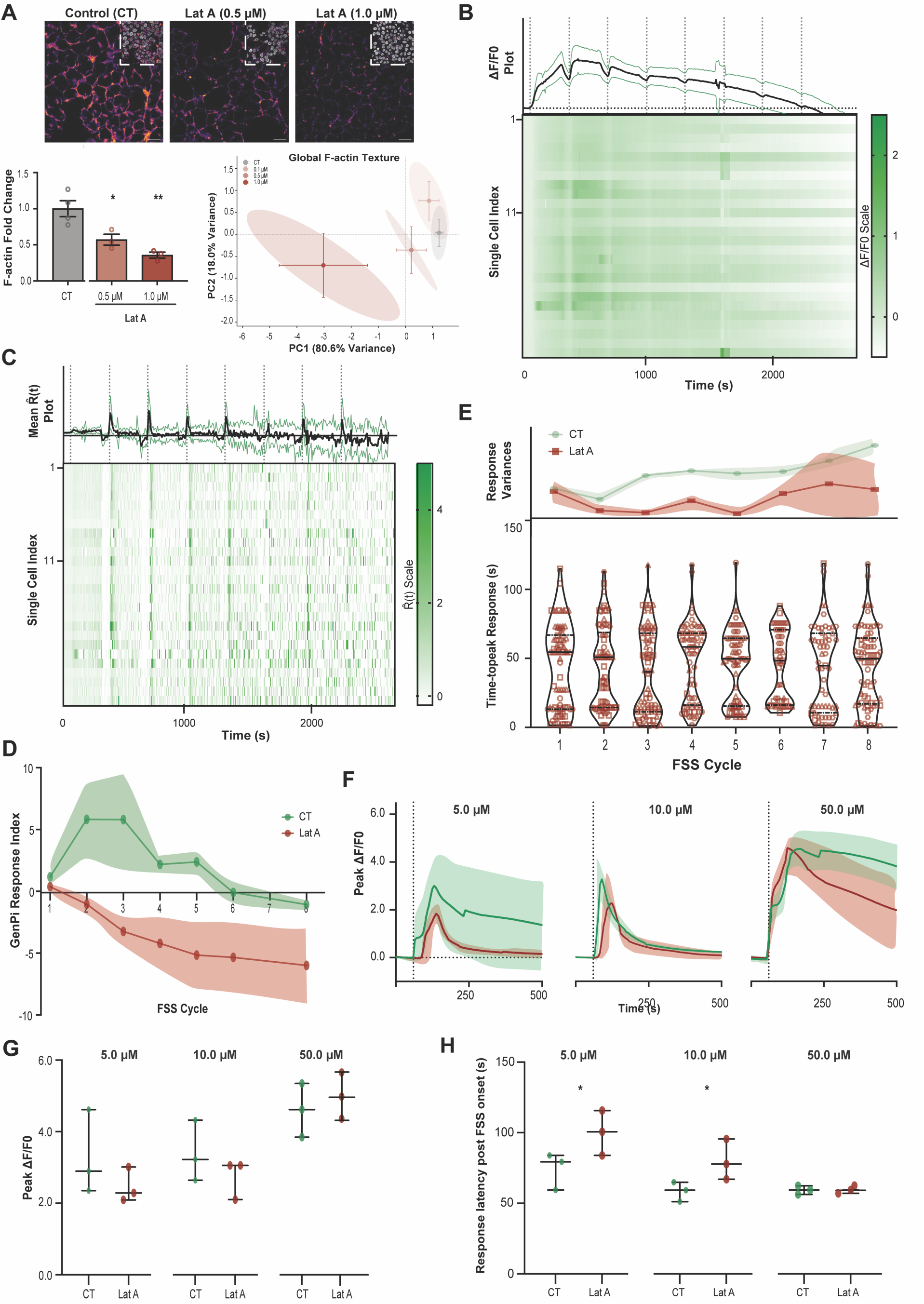
Dose-dependent actin disruption and Piezo1 pharmacological activation following low-dose Latrunculin. **A.** (A) Representative phalloidin-labelled F-actin and DAPI images in control cells and cells exposed to 0.5 or 1.0 µM Latrunculin A (Lat A). Bulk F-actin fluorescence decreased with increasing Lat A concentration, providing a positive-control context for gross cytoskeletal disruption (*N* = 3 biological replicates; One-way analysis of variance (ANOVA); *, *P* = 0.0192 (0.5 µM); **, *P* = 0.0013 (1.0 µM)). The functional mechanotransduction experiments in the main manuscript used the lower 0.1 µM Lat A condition, for which bulk F-actin fluorescence was not significantly reduced (main Fig. 5A). GLCM-based texture analysis quantified five F-actin texture features (contrast, homogeneity, correlation, energy and entropy). Principal component analysis (PCA) was used to visualise multivariate variation in the texture measurements and did not reveal clear separation between CT and 0.1 µM Lat A-treated cells. (B) Population-wide GenEPi responses during repeated 2.5 dynes/cm² fluid shear stress (FSS) stimulation in a representative control (CT) biological replicate. Top: mean ΔF/F₀ trace; bottom: single-cell heatmap. (C) Temporal-derivative representation of population-wide GenEPi activity in control cells during repeated FSS from representative replicate shown in (B). Positive stimulus-associated derivative responses were most pronounced during early stimulation cycles. (D) GenEPi Response Index across independent biological replicates during repeated FSS. Positive values indicate activation-dominant responses and negative values depression-dominant responses. Control cells showed positive stimulus-associated responses during early cycles, whereas low-dose Lat A-treated cells shifted toward negative values. (*N* = 3 biological replicates per condition; Linear mixed-effects model; treatment: *F_1,6_* = 11.01, *P* = 0.016; stimulation cycle: *F_7,42_* = 3.57, *P* = 0.0042; and treatment x stimulation cycle interaction: *F_7,42_* = 2.85, *P* = 0.0157) (E) Response-time dispersion during repeated stimulation. Linear mixed-effects analysis demonstrated significant effects of stimulation cycle (*N* = 3 biological replicates; *F_7_,₄₂* = 5.63, *P* < 0.001) and treatment (*N* = 3 biological replicates; *F₁,₇* = 9.62, *P* = 0.017), whereas the treatment x stimulation cycle interaction was not significant (*N* = 3 biological replicates; *F_7_,₄₂* = 1.76, *P* = 0.132). (F) Representative GenEPi responses following Yoda1 at 5, 10, and 50 µM in control and low-dose Lat A-treated cells. Traces were interpolated onto a common real-time axis without peak alignment; thick lines indicate condition means and shaded regions denote ± SEM. (G) Peak GenEPi responses across independent biological replicates following Yoda1. Peak responses were largely preserved after low-dose Lat A across the tested Yoda1 concentrations (*N* = 3 biological replicates per condition; Two-way analysis of variance (ANOVA); *P* > 0.05 for all tests). (H) Response latency across independent biological replicates following Yoda1. Low-dose Lat A increased response latency at lower Yoda1 concentrations (5 and 10 µM), whereas responses at 50 µM occurred with more similar timing (*N* = 3 biological replicates per condition; Two-way analysis of variance (ANOVA); *, *P* = 0.0121 (5 µM) and 0.0289 (10 µM)).

**Table S1.**
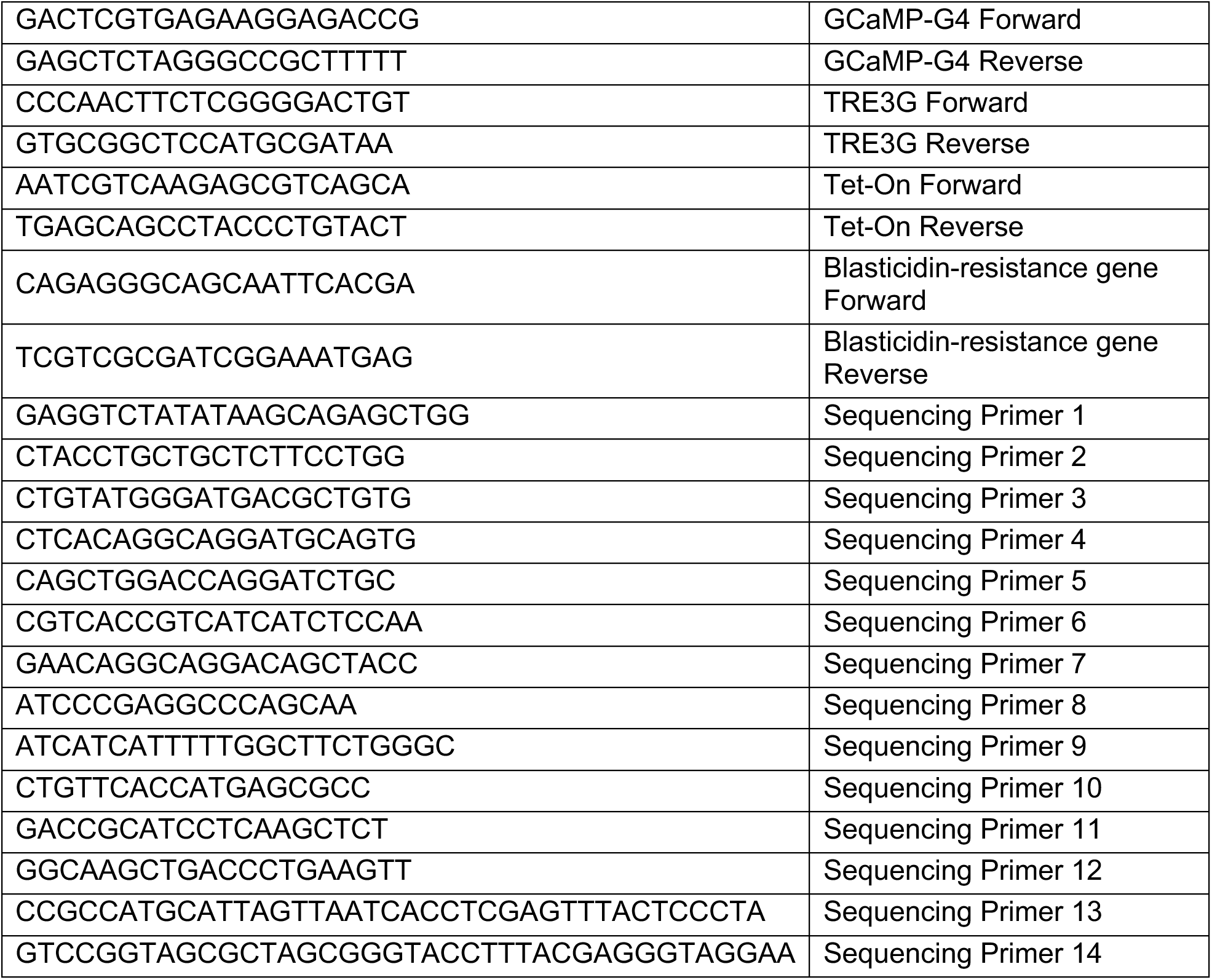
Primer sequences used for cloning and validation. A total of 4 primer pairs were generated using NCBI Primer-Blast designing tool to screen for the four main elements in XLGenEPi construct, namely: (i) GCaMP-G4 sequence, (ii) TRE-promoter sequence, (iii) Tet-On gene sequence, and (iv) blasticidin-resistance gene sequence. An additional 14 primers were generated to screen for both the quality and integrity of the construct from the molecular cloning. All primers detailed in 5’→ 3’ sequence.

**Table S2.** Antibodies and staining reagents used for immunofluorescence.

|  |  |
| --- | --- |
| Anti-Piezo1 antibody | (Proteintech; 15939-1-AP) |
| Donkey anti-rabbit IgG (H+L) Alexa Fluor Plus-488 | (Thermo Fisher, A32790) |
| Alexa Fluor-647 Phalloidin | (Invitrogen, A22287) |
| DAPI | (Thermo Fisher, 62248) |

**Supplementary Movie S1. Low-dose Latrunculin A converts sustained Piezo1-associated responses during repeated FSS into transient responses.**

